# On the inference of complex phylogenetic networks by Markov Chain Monte-Carlo

**DOI:** 10.1101/2020.10.07.329425

**Authors:** Rabier Charles-Elie, Berry Vincent, Glaszmann Jean-Christophe, Pardi Fabio, Scornavacca Celine

**Affiliations:** Institut des Sciences de l’Evolution (ISE-M), Université de Montpellier, CNRS, EPHE, IRD, Montpellier, France; Laboratoire d’Informatique, de Robotique et de Microélectronique de Montpellier (LIRMM), Université de Montpellier, CNRS, Montpellier, France; Institut Montpelliérain Alexander Grothendieck (IMAG), Université de Montpellier, CNRS, Montpellier, France; CIRAD, UMR AGAP, F-34398 Montpellier, France; Amélioration Génétique et Adaptation des Plantes méditerranéennes et tropicales (AGAP), Université de Montpellier, CIRAD, INRAE, Institut Agro, Montpellier, France

## Abstract

For various species, high quality sequences and complete genomes are nowadays available for many individuals. This makes data analysis challenging, as methods need not only to be accurate, but also time efficient given the tremendous amount of data to process. In this article, we introduce an efficient method to infer the evolutionary history of individuals under the multispecies coalescent model in networks (MSNC). Phylogenetic networks are an extension of phylogenetic trees that can contain *reticulate* nodes, which allow to model complex biological events such as horizontal gene transfer, hybridization, introgression and recombination. We present a novel way to compute the likelihood of *biallelic* markers sampled along genomes whose evolution involved such events. This likelihood computation is at the heart of a Bayesian network inference method called SnappNet, as it extends the Snapp method [1] inferring evolutionary trees under the multispecies coalescent model, to networks. SnappNet is available as a package of the well-known beast 2 software.

Recently, the `MCMCBiMarkers` method [2] also extended Snapp to networks. Both methods take biallelic markers as input, rely on the same model of evolution and sample networks in a Bayesian framework, though using different methods for computing priors. However, SnappNet relies on algorithms that are exponentially more time-efficient on non-trivial networks. Using extensive simulations, we compare performances of SnappNet and `MCMCBiMarkers`. We show that both methods enjoy similar abilities to recover simple networks, but SnappNet is more accurate than `MCMCBiMarkers` on more complex network scenarios. Also, on complex networks, SnappNet is found to be extremely faster than `MCMCBiMarkers` in terms of time required for the likelihood computation. We finally illustrate SnappNet performances on a rice data set. SnappNet infers a scenario that is compatible with simpler schemes proposed so far and provides additional understanding of rice evolution.

**Author summary:** Nowadays, to make the best use of the vast amount of genomic data at our disposal, there is a real need for methods able to model complex biological mechanisms such as hybridization and introgression. Understanding such mechanisms can help geneticists to elaborate strategies in crop improvement that may help reducing poverty and dealing with climate change. However, reconstructing such evolution scenarios is challenging. Indeed, the inference of phylogenetic networks, which explicitly model reticulation events such as hybridization and introgression, requires high computational resources. Then, on large data sets, biologists generally deduce reticulation events indirectly using species tree inference tools.

In this context, we present a new Bayesian method, called SnappNet, dedicated to phylogenetic network inference. Our method is competitive in terms of execution speed with respect to its competitors. This speed gain enables us to consider more complex evolution scenarios during Bayesian analyses. When applied to rice genomic data, SnappNet suggested a new evolution scenario, compatible with the existing ones: it posits cAus as the result of an early combination between the Indica and Japonica lineages, followed by a later combination between the cAus and Japonica lineages to derive cBasmati. This accounts for the well-documented wide hybrid compatibility of cAus.

## Introduction

Complete genomes for numerous species in various life domains [3–7], and even for several individuals for some species [8, 9] are nowadays available thanks to next generation sequencing. This flow of data finds applications in various domains such as pathogenecity [10], crop improvement [11], evolutionary genetics [12] or population migration and history [13–15]. Generally, phylogenomic studies use as input thousands to millions of highly conserved genomic fragments, called *genes* (or *loci* when non-coding regions are also concerned). To process such a large amount of data, methods need not only to be accurate, but also time efficient. The availability of numerous genomes at both the intra and inter species levels has been a fertile ground for studies at the interface of population genetics and phylogenetics [16] that aim at estimating the evolutionary history of closely related species. In particular, the well-known coalescent model from population genetics [17] has been extended to the *multispecies coalescent* (MSC) model [18, 19] to handle studies involving populations or individuals from several species. Recent studies on this model show how to incorporate sequence evolution processes from the phylogenetic field into the MSC [1, 20]. As a result, it is now possible to reconstruct evolutionary histories while accounting for both incomplete lineage sorting (ILS) and sequence evolution [21, 22].

For a given locus, ILS leads different individuals in a same population to have different alleles that can trace back to different ancestors. Then, if speciation occurs before that the different alleles are sorted in the population, the locus tree topology can differ from the species history [23]. But incongruence between these trees can also result from biological phenomena that lead distinct species to exchange genetic material. Mechanisms at stake lead a genome to have different parent species – in contrast with the simpler image that depicts a genome as being vertically inherited with modifications from a single ancestral genome. Examples of mechanisms leading to mix genome contents are horizontal gene transfers (present in prokaryotes [24] and eukaryotes [25]), hybridizations (in plants and animals [26–29]), introgressions (e.g. rice [30], citrus [31], sea bass [32]) and recombinations [33]. The latest phenomenon involves species that pair and shuffle related sequences, which allows viruses to produce novel strains evading pre-existing immunity [34]. The evolutionary outcomes of all these *reticulate* events, are largely the same: the production of individuals or lineages originating from the merging of two or more ancestors. As rooted trees are not suited to represent the history of such lineages, they are duly replaced by rooted phylogenetic networks. A rooted phylogenetic network is mainly a directed acyclic graph whose internal nodes can have several children, as in trees, but can also have several parents [35–37]. Various models of phylogenetic network have been proposed over time to explicitly represent reticulate evolution, such as hybridization networks [38] or ancestral recombination graphs [39], along with dozens of inference methods [40, 41].

Model-based methods have been proposed to handle simultaneously ILS and reticulate evolution, which is a desired feature to avoid bias in the inference [42–44]. These methods postulate a probabilistic model of evolution and then estimate its parameters from the data, including the underlying network. The estimation of these parameters such as branch lengths (and hence speciation dates) and population sizes makes them more versatile than combinatorial methods [45]. On the down side, they usually involve high running times as they explore large parameter spaces. Two probabilistic models differentiate regarding the way a locus tree can be embedded within a network. In Kubatko’s model [46, 47], all lineages of a given locus tree coalesce within a single species tree, called parental tree, *displayed* by the network. The model of Yu et al. [48] is more general as, at each reticulation node, a lineage of the locus tree is allowed to descend from a parental ancestor independently of which ancestors provide the other lineages. Works on the latter model extends in various ways the MSC model to consider network-like evolution, giving rise to the *multispecies network coalescent* (MSNC), intensively studied in recent years [2, 41, 44, 49–57]. For this model, Yu et al. have shown how to compute the probability mass function of a non-recombinant locus (*gene*) tree evolving inside a network, given the branch lengths and inheritance probabilities at each reticulation node of the network [49, 51].

This opened the way to infer networks according to the well-known maximum likelihood and Bayesian statistical frameworks.

When the input data consists in multi-locus alignments, a first idea is to decompose the inference process in two steps: first, infer locus trees from their respective alignments, then look for networks that can lead to observe such trees. Following this principle, Yu et al devised a maximum likelihood method [51], then a Bayesian sampling technique [54]. However, using locus trees as a proxy of molecular sequences looses some information contained in the alignments [18] and is subject to gene reconstruction errors. For this reasons, recent work considers jointly estimating the locus trees and the underlying network. This brings the extra advantage that better locus trees are likely to be obtained [58], but running time may become prohibitive already for inferences on few species. Wen et al in the PhyloNet software [55] and Zhang et al. with the SpeciesNetwork method [56] both proposed Bayesian methods following this principle.

Though a number of trees for a same locus are considered during such inference processes, they are still considered one at a time, which may lead to a precision loss (and a time loss) compared to an inference process that would consider all possible trees for a given locus at once. When data consists in a set of *biallelic* markers (e.g., SNPs), the ground-breaking work of Bryant et al. [1] allows to compute likelihoods while integrating over all gene trees, under the MSC model (*i*.*e*., when representing the history as a tree). This work was recently extended to the MSNC context by Zhu et al [2].

In this paper, we present a novel way to compute the probability of biallelic markers, given a network. This likelihood computation is at the heart of a Bayesian network inference method we called SnappNet, as it extends the Snapp method [1] to networks. SnappNet is available at https://github.com/rabier/MySnappNet and distributed as a package of the well-known beast 2 software [59, 60]. This package partly relies on code from Snapp [1] to handle sequence evolution and on code from SpeciesNetwork [56] to modify the network during the MCMC as well as to compute network priors.

Our approach differs from that of Zhang et al. [56] in that SnappNet takes a matrix of biallelic markers as input while SpeciesNetwork expects a set of alignments. Thus, the considered substitution models differ as we consider only two states (alleles) while SpeciesNetwork deals with nucleotides. The computational approaches also differ as our MCMC integrates over all locus trees for each sampled network, while SpeciesNetwork jointly samples networks and gene trees. Though summarizing the alignments by gene trees might be less flexible, this allows SpeciesNetwork to provide embeddings of the gene trees into the sampled networks, while in our approach this needs to be done in a complementary step after running SnappNet. However, managing the embeddings can also lead to computational issues as Zhang et al. report, since a topological change for the network usually requires a recomputation of the embeddings for all gene trees [56].

The SnappNet method we present here is much closer to the `MCMCBiMarkers` method of Zhu et al. [2], which also extends the Snapp method [1] to network inference. Both methods take biallelic markers as input, rely on the same model of evolution and also both sample networks in a Bayesian framework. However, the methods differ in two important points: the way the Bayesian inference is conducted and, most importantly, in the way likelihoods are computed. The result section shows that this often leads to tremendous differences in running time, but also to differences in convergence.

We note here that reducing running times of model-based methods can also be done by approximating likelihoods, as done by *pseudo-likelihood* methods: the network likelihood is computed for subparts of its topology, these values being then assembled to approximate the likelihood of the full network. A decomposition of the network into rooted networks on three taxa (trinets) is proposed in the PhyloNet software [52, 61] and one into semi-directed networks on four taxa in the SNaQ method of the PhyloNetwork package [53]. Since pseudo-likelihood methods are approximate heuristics to compute a likelihood, they are usually much faster than full likelihood methods and can handle large genomic data sets. On the downside, these methods face, more often that the full-likelihood methods, serious identifiability problems since some networks can simply not be recovered from topological substructures such as rooted triples, quartets or even embedded trees [52, 53, 62]. This is why we decided to focus on the *exact* computation of the *full* likelihood.

In the following, we first detail the mathematical model considered, then explain the SnappNet method, before illustrating its performances on simulated and real data.

## Materials and methods

### Input data

SnappNet considers as input data a matrix *D* containing an alignment of *m* biallelic markers sampled from a number of individuals. Each individual belongs to a given species. These species are in a 1-to-1 correspondence with the leaves of an unknown phylogenetic network, which is the main parameter that we wish to estimate. The markers can be SNPs or random sites sampled from chromosomes, including invariant sites. All markers are considered to be independent, so a certain distance must be preserved between genomic locations included in the matrix. We identify the two alleles with the colors red and green.

Each column *D*_*i*_ of the alignment corresponds to a different marker. The only information that is relevant to SnappNet’s computations are the numbers of red and green alleles observed in *D*_*i*_ for the individuals of a given species. This implies that unphased data can be analyzed with SnappNet, as long as it is translated in the input format expected by the software.

### Mathematical model

In this paper, we refer to phylogenetic networks as directed acyclic graphs with branches oriented as the time flows, see Figure 1. At their extremities, networks have a single node with no incoming branch and a single outgoing branch —the *origin*— and a number of nodes with a single incoming branch and no outgoing branches —the *leaves*. All other nodes either have a single incoming branch and two outgoing branches —the *tree* nodes— or two incoming branches and a single outgoing branch —the *reticulation* nodes. Tree nodes and reticulation nodes represent speciations and hybridization events, respectively. For consistency with Zhang et al. [56], the immediate descendant of the origin – that is, the tree node representing the first speciation in the network – is called the *root*.

**Fig 1.**
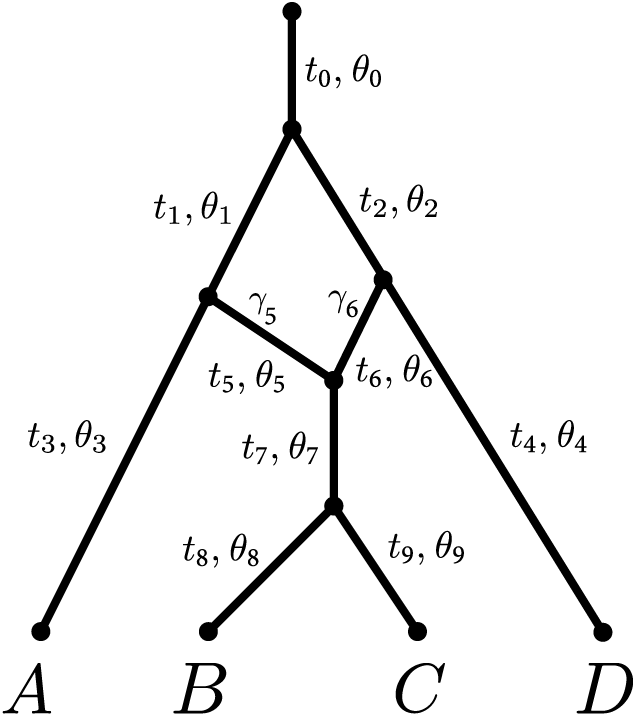
Example of a phylogenetic network. The top node represents the origin and its child node is called the root of the network. Time flows from the origin node to the leaves (here *A, B, C, D*) so branches are directed from the top to the leaves. Each branch *x* is associated to a length *t*_*x*_, and to a population size *θ*_*x*_. Additionally, branches *x* on top of a reticulation node have an inheritance probability *γ*_*x*_ representing their probability to have contributed to any individual at the top of the branch just below.

Each branch *x* in the network represents a population, and is associated to two parameters: a scaled population size *θ*_*x*_ and a branch length *t*_*x*_. Any branch *x* on top of a reticulation node *h* is further associated with a probability *γ*_*x*_ *∈* (0, 1), under the constraint that the probabilities of the two parent branches of *h* sum to 1. These probabilities are called *inheritance probabilities*. All these parameters have a role in determining how gene trees are generated by the model, and how markers evolve along these gene trees, as described in the next two subsections, respectively.

### Gene tree model

Gene trees are obtained according to the MSNC model. The process starts at the leaves of the network, where a given number of lineages is sampled for each leaf, each lineage going backwards in time, until all lineages coalesce. Along the way, this process determines a gene tree whose branch lengths are each determined as the amount of time between two coalescences affecting a single lineage. In what follows, “times” —and therefore branch lengths— are always measured in terms of expected number of mutations per site.

Within each branch *x* of the network, the model applies a standard coalescent process governed by *θ*_*x*_. In detail, any two lineages within *x* coalesce at rate 2*/θ*_*x*_, meaning that the first coalescent time among *k* lineages follows an exponential distribution *ε* (*k*(*k –* 1)*/θ*_*x*_), since the coalescence of each combination of 2 lineages is equiprobable. Naturally, if the waiting time to coalescence exceeds the branch length *t*_*x*_, the lineages are passed to the network branch(es) above *x* without coalescence. If there are two such branches *y, z* (i.e., the origin of *x* is a reticulation node), then each lineage that has arrived at the top of branch *x* chooses independently whether it goes to *y* or *z* with probabilities *γ*_*y*_ and *γ*_*z*_ = 1 – *γ*_*y*_, respectively [48]. The process terminates when all lineages have coalesced and only one ancestral lineage remains.

### Mutation model

As is customary for unlinked loci, we assume that the data is generated by a different gene tree for each biallelic marker. The evolution of a marker along the branches of this gene tree follows a two-states asymmetric continuous-time Markov model, scaled so as to ensure that 1 mutation is expected per time unit. This is the same model as Bryant et al. [1]. For completeness, we describe this mutation model below.

We represent the two alleles by red and green colors. Let *u* and *v* denote the instantaneous rates of mutating from red to green, and from green to red, respectively. Then, for a single lineage, ℙ(red at *t* + Δ*t* | green at *t*) = *v*Δ*t* + *o*(Δ*t*), and ℙ (green at *t* + Δ*t* | red at *t*) = *u*Δ*t* + *o*(Δ*t*), where *o*(Δ*t*) is negligible when Δ*t* tends to zero. The stationary distribution for the allele at the root of the gene tree is green with probability *u/*(*u* + *v*) and red with probability *v/*(*u* + *v*). Under this model, the expected number of mutations per time unit is 2*uv/*(*u* + *v*). In order to measure time (branch lengths) in terms of expected mutations per site (i.e. genetic distance), we impose the constraint 2*uv/*(*u* + *v*) = 1 as in [1]. When *u* and *v* are set to 1, the model is also known as the Haldane model [63] or the Cavender-Farris-Neyman model [64].

## Bayesian framework

### Posterior distribution

Let *D*_*i*_ be the data for the *i*-th marker. The posterior distribution of the phylogenetic network Ψ can be expressed as:

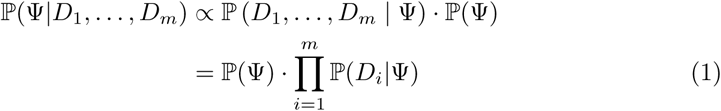

where *∝* means “is proportional to”, and where ℙ (*D*_1_, *…, D*_*m*_ | Ψ) and ℙ(Ψ) refer to the likelihood and the network prior, respectively.

Equation (1) —which relies on the independence of the data at different markers— allows us to compute a quantity proportional to the posterior by only using the prior of Ψ and the likelihoods of Ψ with respect to each marker, that is ℙ(*D*_*i*_ | Ψ). While we could approximate ℙ(*D*_*i*_ |Ψ) by sampling gene trees from the distribution determined by the species network, this is time-consuming and not necessary. Similarly to the work by Bryant et al. [1] for inferring phylogenetic *trees*, we show below that ℙ(*D*_*i*_ |Ψ) can be computed for *networks* using dynamic programming.

SnappNet samples networks from their posterior distribution by using Markov chain Monte-Carlo (MCMC) based on Equation (1).

### Priors

Before describing the network prior, let us recall the network components: the topology, the branch lengths, the inheritance probabilities and the populations sizes. In this context, we used the birth hybridization process of Zhang et al. [56] to model the network topology and its branch lengths. This process depends on the speciation rate *λ*, on the hybridization rate *ν* and on the time of origin *τ*_0_. Hyperpriors are imposed onto these parameters. An exponential distribution is used for the hyperparameters *d* := *λ – ν* and *τ*_0_. The hyperparameter *r* := *ν/λ* is assigned a Beta distribution. We refer to [56] for more details. The inheritance probabilities are modeled according to a uniform distribution. Moreover, like Snapp, SnappNet considers independent and identically distributed Gamma distributions as priors on population sizes *θ*_*x*_ associated to each network branch. This prior on each population size induces a prior on the corresponding coalescence rate (see [1] and Snapp’s code). Last, as in Snapp, the user can specify fixed values for the *u* and *v* rates, or impose a prior for these rates and let them be sampled within the MCMC.

### Partial likelihoods

In the next section we describe a few recursive formulae that we use to calculate the likelihood ℙ(*D*_*i*_ |Ψ) using a dynamic programming algorithm. Here we introduce the notation that allows us to define the quantities involved in our computations. Unless otherwise stated, notations that follow are relative to the *i*th biallelic marker. To keep the notations light, the dependence on *i* is not explicit.

Given a branch *x*, we denote by 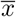 and *<u>x</u>* the top and bottom of that branch. We call 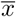 and *<u>x</u> population interfaces*. 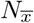 and *N*_*<u>x</u>*_ are random variables denoting the number of gene tree lineages at the top and at the bottom of *x*, respectively. Similarly, 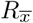 and *R*_*<u>x</u>*_ denote the number of red lineages at the top and bottom of *x*, respectively.

For simplicity, when *x* is a branch incident to a leaf, we identify *<u>x</u>* with that leaf. Two quantities that are known about each leaf are *r*_*<u>x</u>*_ and *n*_*<u>x</u>*_, which denote the number of red lineages sampled at *<u>x</u>* and the total number of lineages sampled at *<u>x</u>*, respectively. Note that *N*_*<u>x</u>*_ is non-random: indeed, it must necessarily equal *n*_*<u>x</u>*_, which is determined by the number of individuals sampled from that species. On the other hand, the model we adopt determines a distribution for the *R*<u>_*x*_</u>. The probability of the observed values *r*<u>_*x*_</u> for these random variables equals ℙ(*D*_*i*_|Ψ).

Now let **x** be an ordered collection (i.e. a vector) of population interfaces. We use **n**_**x**_ (or **r**_**x**_) to denote a vector of non-negative integers in a 1-to-1 correspondence with the elements of **x**. Then *N*_**x**_ = **n**_**x**_ is a shorthand for the equations expressing that the numbers of lineages in **n**_**x**_ are observed at their respective interfaces in **x**. For example, if 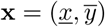 and **n**_**x**_ = (*m, n*), then *N*_**x**_ = **n**_**x**_ is a shorthand for *N*_*<u>x</u>*_ = *m*, 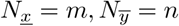. We use *R*_**x**_ = **r**_**x**_ analogously to express the observation of the numbers of red lineages in **r**_**x**_ at **x**.

In order to calculate the likelihood ℙ(*D*_*i*_ | Ψ), we subdivide the problem into that of calculating quantities that are analogous to partial likelihoods. Given a vector of population interfaces **x**, let **L**(**x**) denote a vector containing the leaves that descend from any element of **x**, and let **r**_**L**(**x**)_ be the vector containing the numbers of red lineages *r*<u>_*x*_</u> observed at each leaf *<u>x</u>* in **L**(**x**). Then we define:

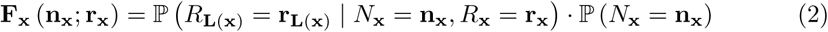

These quantities are generalizations of similar quantities defined by Bryant et al. [1]. We will call them partial likelihoods, although, as noted by these authors, strictly speaking this is an abuse of language.

### Computing partial likelihoods: the rules

Here we show a set of rules that can be applied to compute partial likelihoods in a recursive way. Derivations and detailed proofs of the correctness of these rules can be found in the Supplementary Materials.

We use the following conventions. In all the rules that follow, vectors of population interfaces **x, y, z** are allowed to be empty. The comma operator is used to concatenate vectors or append new elements at the end of vectors, for example, if **a** = (*a*_1_, *a*_2_, *…, a*_*k*_) and **b** = (*b*_1_, *b*_2_, *…, b*_*h*_), then **a, b** = (*a*_1_, *…, a*_*k*_, *b*_1_, *…, b*_*h*_) and **a**, *c* = (*a*_1_, *a*_2_, *…, a*_*k*_, *c*). Trivially, if **a** is empty, then **a, b** = **b** and **a**, *c* = (*c*). A vector **x** of *incomparable* population interfaces is such that no two elements of **x** are equal, nor one is descendant of the other in the network. Finally, for any branch *x*, let *m*_*x*_ denote the number of lineages sampled in the descendant leaves of *x*.

**Rule 0:** Let *x* be a branch incident to a leaf. Then,

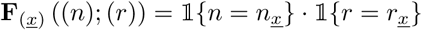

**Rule 1:** Let **x**, *<u>x</u>* be a vector of incomparable population interfaces. Then,

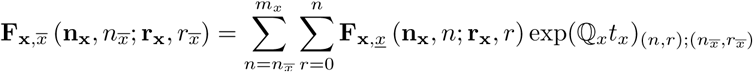

where *t*_*x*_ denotes the length of branch *x*, and ℚ_*x*_ is the rate matrix defined by Bryant et al. [1, p. 1922] that accounts for both coalescence and mutation (see also the Supplementary Materials).

**Rule 2:** Let 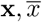 and 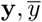 be two vectors of incomparable population interfaces, such that 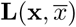 and 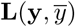 have no leaf in common. Let *x, y* be the immediate descendants of branch *z*, as in Fig. 2. Then,

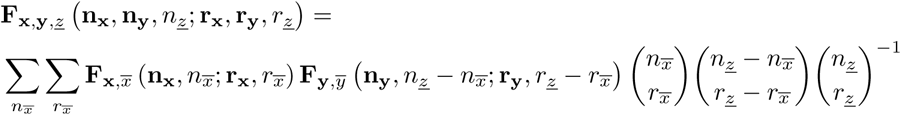

The ranges of 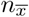 and 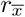 in the summation terms are defined by 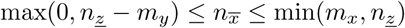 and 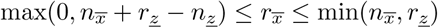.

**Fig 2.**
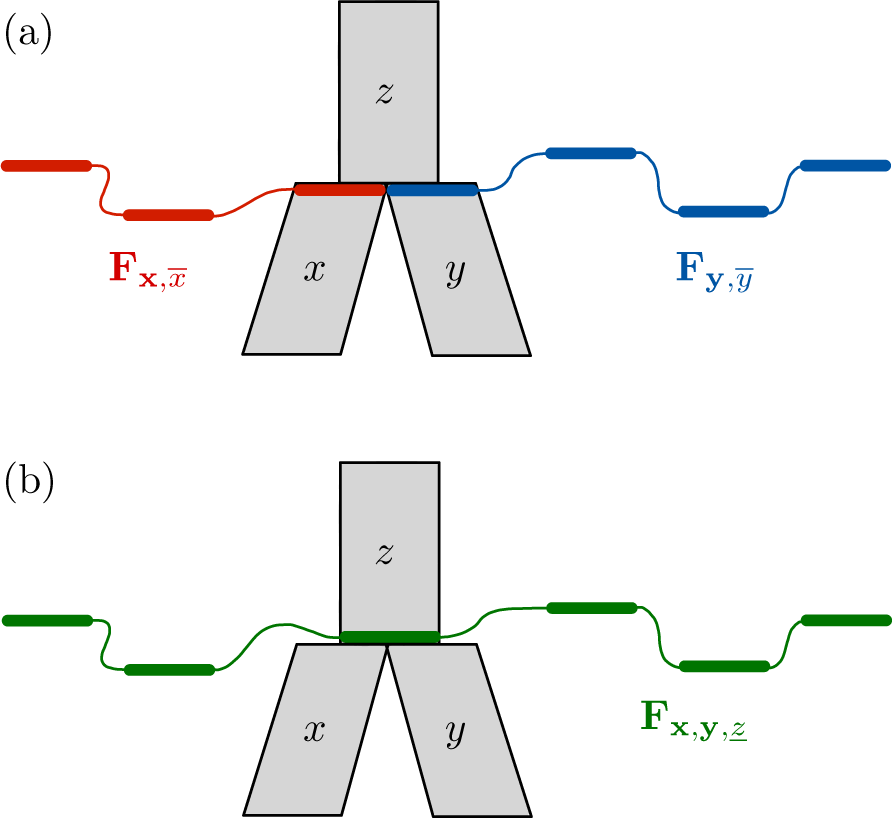
Illustration of Rule 2. Given (a) the partial likelihoods for the 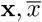 (red) vector of population interfaces and the partial likelihoods for the 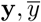 (blue) vector of population interfaces, Rule 2 allows us to compute the partial likelihoods for the (green) vector **x, y**, *<u>z</u>* (b).

**Rule 3:** Let 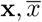 be a vector of incomparable population interfaces, such that branch *x*’s top node is a reticulation node. Let *y, z* be the branches immediately ancestral to *x*, as in Fig. 3. Then,

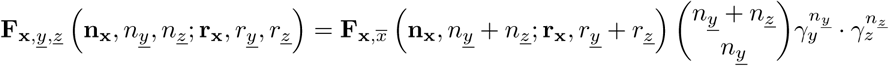

**Rule 4:** Let 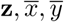 be a vector of incomparable population interfaces, and let *x, y* be immediate descendants of branch *z*, as in Fig. 4. Then,

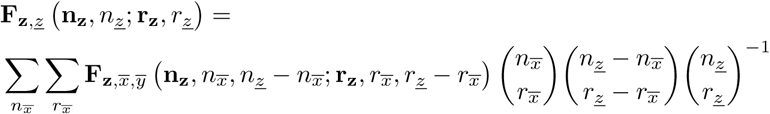

The ranges of 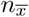 and 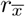 in the sums are the same as those in Rule 2.

Repeated application of the rules above, performed by an algorithm described in the next subsection, leads eventually to the partial likelihoods for *<u>ρ</u>*, the population interface immediately above the root of the network (i.e, *ρ* is the branch linking the origin to the root). From these partial likelihoods, the full likelihood ℙ(*D*_*i*_|Ψ) is computed as follows:

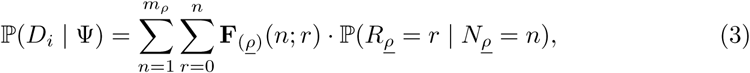

where the conditional probabilities ℙ(*R*_*<u>ρ</u>*_ = *r* | *N*<u>_*ρ*_</u> = *n*) are obtained as described by Bryant et al. [1]. Note that the length of branch *ρ* does not play any role in the computation of the likelihood, so it is not identifiable.

**Fig 3.**
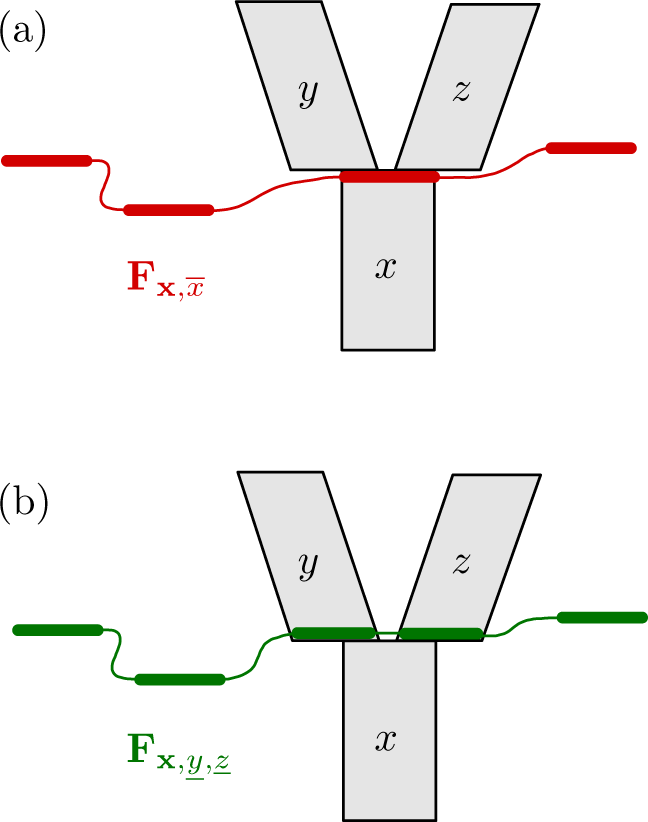
Illustration of Rule 3. Given (a) the partial likelihoods for the 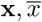 (red) vector of population interfaces, Rule 3 allows us to compute the partial likelihoods for the (green) vector **x**, *<u>y</u>, <u>z</u>* (b).

### Likelihood computation

We now describe the algorithm that allows SnappNet to derive the full likelihood ℙ(*D*_*i*_ |Ψ) using the rules introduced above. We refer to the Supplementary Materials for detailed pseudocode.

The central ingredient of this algorithm are the partial likelihoods for a vector of population interfaces **x** (here referred to as VPI), which are stored in a matrix with potentially high dimension, denoted **F**_**x**_. We say that a VPI **x** is *active* at some point during the execution of the algorithm, if: (1) **F**_**x**_ has been computed by the algorithm, (2) **F**_**x**_ has not yet been used to compute the partial likelihoods for another VPI. To reduce memory usage, we only store **F**_**x**_ for active VPIs.

In a nutshell, the algorithm traverses each node in the network following a topological sort [65], that is, in an order ensuring that a node is only traversed after all its descendants have been traversed. Every node traversal involves deriving the partial likelihoods of a newly active VPI from those of at most two VPIs that, as a result, become inactive. Eventually, the root of the network is traversed, at which point the only active VPI is (*<u>ρ</u>*) and the full likelihood of the network is computed from **F**_(*<u>ρ</u>*)_ using Equation (3).

In more detail, a node is ready to be traversed when all its child nodes have been traversed. At the beginning, only leaves can be traversed and their partial likelihoods **F**_(*<u>x</u>*)_ are obtained by application of Rule 0, followed by Rule 1 to obtain 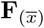. Every subsequent traversal of a node *d* entails application of one rule among Rules 2, 3 or 4, depending on whether *d* is a tree node and on whether the branch(es) topped by *d* correspond to more than one VPI (see Figs. 2–4). The selected rule computes **F**_**x**_ for a newly active VPI **x**. This is then followed by application of Rule 1 to replace every occurrence of any population interface *<u>x</u>* in **x** with 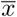.

It is helpful to note that at any moment, the set of active VPIs forms a frontier separating the nodes that have already been traversed, from those that have not yet been traversed (i.e., if branch *x* = (*d, e*) with *d* not traversed and *e* traversed, then there must be an active VPI with *<u>x</u>* or 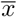 among its population interfaces). Any node that lies immediately above this frontier can be the next one to be traversed. Thus, there is some latitude in the choice of the complete order in which nodes are traversed. Different orders will lead to different VPIs being activated by the algorithm, which in turn will lead to different running times. In fact, running times are largely determined by the sizes of the VPIs encountered. This point is explored further in the next section.

**Fig 4.**
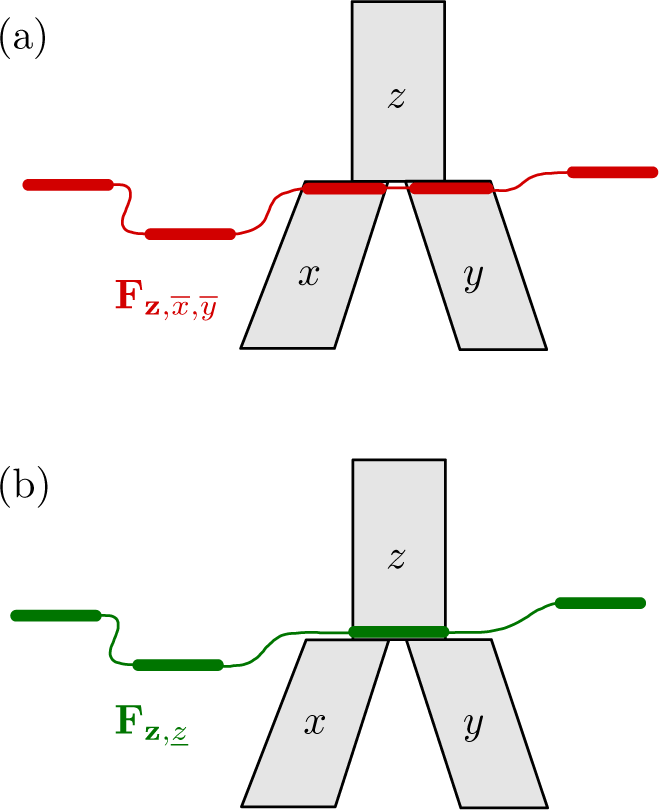
Illustration of Rule 4. Given (a) the partial likelihoods for the 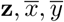 (red) vector of population interfaces, Rule 4 allows us to compute the partial likelihoods for the (green) vector **z**, *<u>z</u>* (b).

### Time complexity of computing the likelihood

Our approach improves the running times by several orders of magnitude with respect to previous work to compute the likelihood of a network for biallelic marker data [2]. This is clearly apparent for some experiments detailed in the Results section, but it can also be understood by comparing computational complexities.

Here, let *n* be the total number of individuals sampled, and let *s* denote the size of the species network Ψ (i.e. its number of branches or its number of nodes). Let us first examine the running time to process one node in Ψ. For any of Rules 0-4, let *K* be the number of population interfaces in the VPI for which partial likelihoods are being computed, that is, *K* is the number of elements of 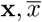 for Rule 1, that of **x, y**, *<u>z</u>* for Rule 2, and so on. These partial likelihoods are stored in a 2*K*-dimensional matrix, with *O*(*n*^2*K*^) elements. Each rule specifies how to compute an element of this matrix in at most *O*(*n*^2^) operations (in fact rules 0 and 3 only require *O*(1) operations). Thus, any node in the network can be processed in *O*(*n*^2*K*+2^) time.

Since the running time of any other step – i.e. computing Equation (3), and exp(ℚ_*x*_*t*_*x*_) – is dominated by these terms, the total running time is *O*(*sn*^2*K*+2^), where 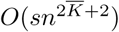 is the maximum number of population interfaces in a VPI activated by the given traversal.

**Fig 5.**
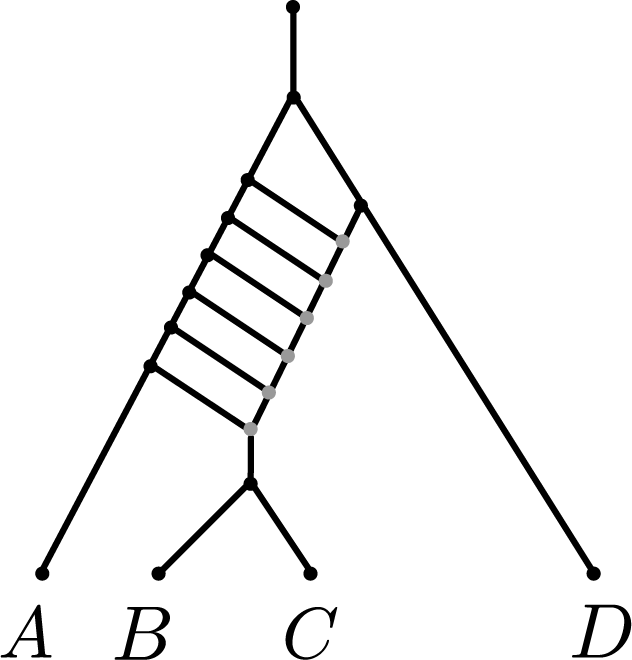
Example of a phylogenetic network where the level 𝓁 is equal to 6 (the reticulation nodes are depicted in grey), while 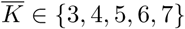, depending on the traversal algorithm (not shown). A traversal ensuring that 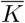 remains close to the lower end of this interval (the scanwidth of the network) will be several orders of magnitude faster than algorithms whose complexity depends exponentially on 𝓁. Increasing the number of reticulation nodes while keeping a “ladder” topology as above can make 𝓁 arbitrarily large, while the scanwidth remains constant. This topology may seem odd but it is intended as the backbone of a more complex and realistic network with subtrees hanging from the different internal edges of the ladder, in which case the complexity issue remains.

Let us now compare this to the complexity of the likelihood computations described by Zhu et al. [2]. Processing a node *d* of the network in their algorithm involves at most *O*(*n*^4*rd*^ +4) time, where *r*_*d*_ is the number of reticulation nodes which descend from *d*, and for which there exists a path from *d* that does not pass via a *lowest articulation node* (see definitions in Zhu et al. [2]). In the Supplementary Materials, we show that this entails a total running time of *O*(*sn*^4𝓁+4^), where 𝓁 is the *level* of the network [35, 66].

Thus, the improvement in running times with respect to the algorithm by Zhu et al. [2] relies on the fact that 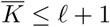. One way of seeing this is to remark that, for any traversal of the network, 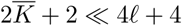. We refer to the Supplementary Materials for a proof of this result. Assuming that 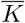 and 𝓁 are close, this would imply that the exponent of *n* in the worst-case time complexity is roughly halved with respect to Zhu et al. [2]. However, 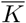 is potentially much smaller than the level 𝓁, as depicted in Figure 5.

We call the minimum value of 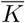 over all possible traversals of the network the *scanwidth* of the network [67]. The current implementation of SnappNet chooses an arbitrary traversal of the network, but research is ongoing to further lower running times by relying on more involved traversal algorithms producing VPIs with sizes closer to the scanwidth [67].

### MCMC operators

SnappNet incorporates the MCMC operators of SpeciesNetwork [56] to move through the network space, and also benefits from operators specific to the mathematical model behind Snapp [1] (e.g. population sizes, mutation rates …). Snapp’s operators on population sizes have been adapted since networks contain more edges than species trees. Moreover, operators on gene trees from [56] have been discarded. As a consequence, SnappNet relies on 16 MCMC operators, described in SnappNet’s manual (https://github.com/rabier/MySnappNet).

## Simulation study

### Simulated data

We implemented a simulator called SimSnappNet, an extension to networks of the SimSnapp software [1]. SimSnappNet handles the MSNC model whereas SimSnapp relies on the MSC model. SimSnappNet is available at https://github.com/rabier/SimSnappNet. In all simulations, we considered a given phylogenetic network, and a gene tree was simulated inside the network, according to the MSNC model. Next, a Markov process was generated along the gene tree branches, in order to simulate the evolution of marker. Note that markers at different sites rely on different gene trees. In all cases, we set the *u* and *v* rates to 1. Then, SnappNet‘s constraint 2*uv/*(*u* + *v*) = 1 is fulfilled. Moreover, we used the same *θ* = 0.005 value, for all network branches. In that sense, our configuration differs slightly from the one of [2]. Recall that these authors considered *θ* = 0.006 for external branches and *θ* = 0.005 for internal branches. Indeed, since SnappNet considers the same prior distribution, Γ(*α, β*), for all *θ*’s, we found more appropriate to generate data under SnappNet‘s assumptions.

Three numbers of markers were studied: either a) 1,000, b) 10,000 or c) 100,000 biallelic sites were generated. Constant sites were not discarded since SnappNet‘s mathematical formulas rely on random markers. However, when the analysis relied only on polymorphic sites, the gene tree and the associated marker were regenerated until having a polymorphic site. We considered 20 replicates for each simulation set up.

### Phylogenetic networks studied

We studied the three phylogenetic networks, presented in Figure 6. Networks A and B are taken from [2]. Network A is the least complex, being *level-1* (i.e. having one reticulation), while network B has two reticulations, located in distinct part of its topology. We also studied network C, a level-2 network where the reticulations are on top of one another, hence have a combined influence on some leaves. In order to fully describe these networks, let us give their extended newick representation [68].

**Fig 6.**
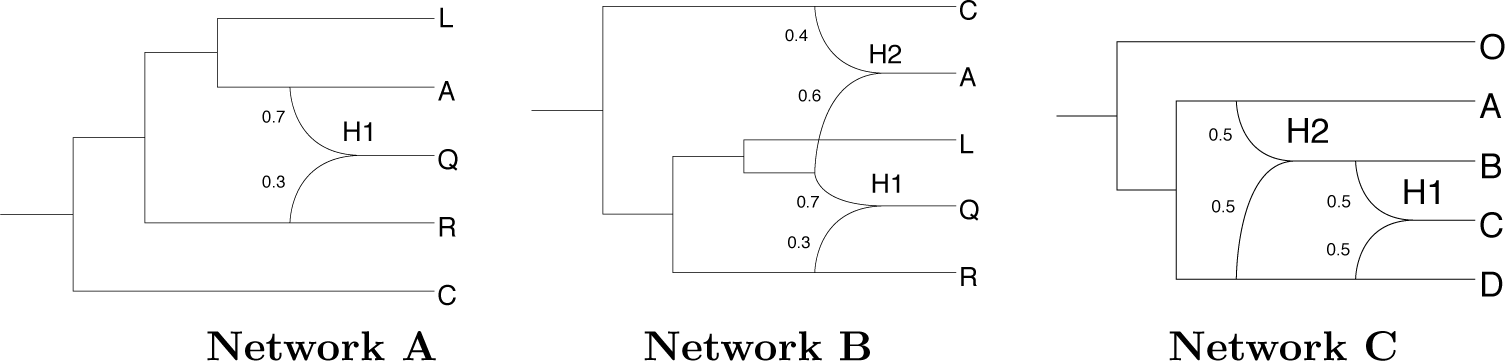
The three phylogenetic networks used for simulating data. Networks A and B are taken from [2]. Branch lengths are measured in units of expected number of mutations per site (i.e. substitutions per site). Displayed values represent inheritance probabilities.

<u>Network A:</u>

~~~
((C:0.08,((R:0.007,(Q:0.004)#H1:0.003):0.035,((A:0.006,#H1:0.002):0.016,
L:0.022):0.02):0.038):0);
~~~

<u>Network B:</u>

~~~
((((R:0.014,(Q:0.004)#H1:0.01):0.028,(((A:0.003)#H2:0.003,#H1:0.002) :0.016,L:0.022):0.02):0.038,(C:0.005,#H2:0.002):0.075):0);
~~~

<u>Network C:</u>

~~~
((O:0.08,((A:0.012,((B:0.002,(C:0.001)#H1:0.001):0.002)#H2:0.008):0.038, ((D:0.003,#H1:0.002):0.017,#H2:0.016):0.03):0.03):0);
~~~

We also focused on networks C(3) and C(4) represented in Figure 7 that are variants from network C. Network C(k), that contains k reticulation nodes, is obtained by splitting species *C* in *k –* 1 subspecies, named *C*_1_, *C*_2_, …, *C*_*k-*1_, and in adding reticulations between them (see Figure 7 for more details).

**Fig 7.**
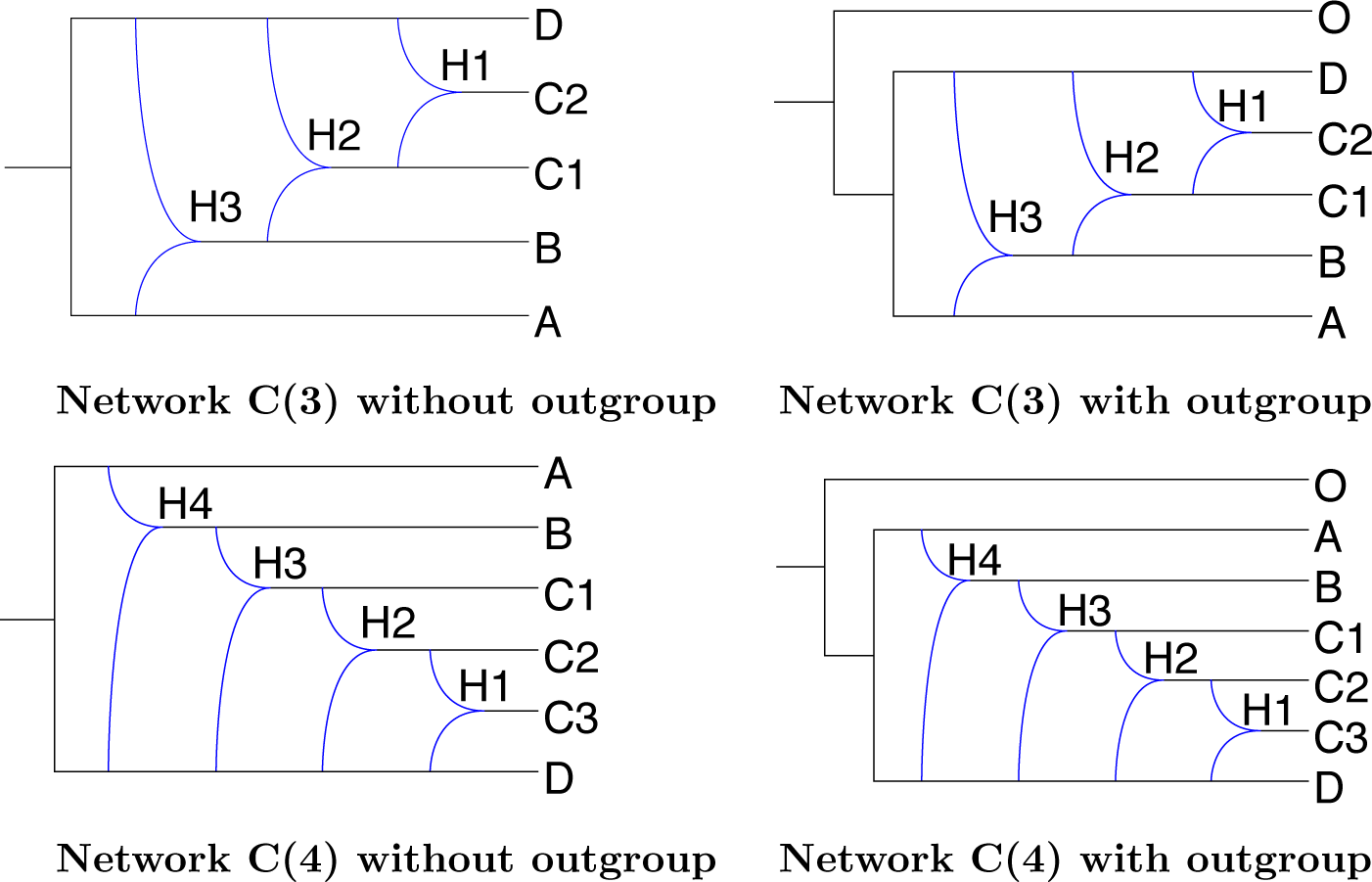
The networks from the C family, with either 3 or 4 reticulation nodes, and with or without outgroup O.

### Bayesian analysis

For studying networks A and B, we used the following species tree as a starting point of the MCMC analysis:

~~~
(((C:0.05,R:0.05):0.05,((A:0.05,L:0.05):0.025,Q:0.075):0.025):0);
~~~

In the same way, for network C, we used the following species tree:

~~~
(((O:0.05,A:0.05):0.05,((C:0.05,D:0.05):0.025,B:0.075):0.025):0);
~~~

As priors on population sizes, we considered *θ ∼* Γ(1, 200) for all branches. Since simulated data were generated by setting *θ* = 0.005, the expected value of this prior distribution is exactly matching the true value (𝔼 (*θ*) = 0.005). For calibrating the network prior, we chose the same distributions as suggested in [56]: *d ∼ε* (0.1), *r ∼* Beta(1, 1), *τ*_0_ *∼ ε* (10). This network prior enables to explore a large network space, while imposing more weights on networks with 1 or 2 reticulations (see Figure 1 of the Supplementary Material). Recall that network A is a 1-reticulation network, whereas networks B and C are 2-reticulation networks. However, in order to limit the computational burden for network C (and for estimating continuous parameters on network A), we modified slightly the prior by bounding the number of reticulations to 2. Last, on network B, an extra analysis was performed by bounding the number of reticulations to 3 in order to compare SnappNet’s results with those obtained by `MCMCBiMarkers` [2]. We refer to Figures 2 and 3 of the Supplementary Material for illustrations of the “bounded” prior.

### MCMC convergence

To track the behaviour of the Bayesian algorithm, we used the Effective Sample Size (ESS) criterion [69]. We assumed that the MCMC convergence was reached and that enough “independent” observations were sampled, when the ESS was greater than 200 (see https://beast.community/ess_tutorial). This threshold is commonly adopted in the MCMC community. The first 10% samples were discarded as burn-in and the ESS was computed on the remaining observations, thanks to the Tracer software [70]. When we could not reach an ESS of 200, the ESS value is specified in the text.

### Accuracy of SnappNet

In order to evaluate SnappNet‘s ability to recover the true network topology, the following accuracy criterion was used. For each replicate, after having discarded the burnin, we computed the ratio, number of observations matching the true topology divided by MCMC chain length. Recall that in our study, the chain length depends on the ESS criterion. Next, the average ratio over the different replicates was computed.

### Real data study on rice

The genomic data are an extract of [71]. We focused on 24 representative cultivars (see Table 3 and Figure 5 in Supplementary Material) spanning the four main rice sub-populations: Indica, Japonica, cAus and cBasmati. We built two data sets of 12 cultivars, each one containing 3 varieties for Indica, Japonica and cBasmati, and 2 varieties for cAus. Data set 1 spans 7 countries from India to the Philippines. Varieties included in Data set 2 come from 8 countries spanning from Pakistan to Indonesia. For each of the 12 chromosomes, we sampled 1k SNPs having only homozygous alleles. Following recommendations of [1], the SNPs were chosen for each of the 12 chromosomes to be as separated as possible from one another to avoid linkage between loci, though [2] has shown this kind of analysis is quite robust to this bias. The concatenation of these SNPs lead to two 12k whole-genome SNP data sets on the selected rice varieties. Last, in order to evaluate the influence of character sampling, a second sampling of 12k SNPs along the whole genome was also considered, for each set of 12 cultivars.

The likelihoods of 10 chosen networks were computed, thanks to SnappNet. Each network represents a different rice evolution scenario. We selected six plausible scenarios and four unlikely scenarios on the bases of the conformity with the main features of the genetic structure of the species: a closer resemblance between Indica and cAus on one side of the structure and the same between Japonica and cBasmati on the other. The former include the scenarios that have most recently been put forward.

The likelihood optimization was based on 9 operators among the 16 original operators presented in [56]. Indeed, in its current version, SnappNet incorporates the 5 topological operators of [56] to explore the space of network topologies, and also 2 operators associated to the network hyperparameters *r* and *d* involved in the prior. Besides, contrary to the MCMC analysis, new parameter values were always accepted as soon as they led to a likelihood increase. The optimization was assumed to be completed when the relative difference (between two states distant from 100 iterations) was found below the threshold 10^*-*6^. Another optimization was also studied, which can be viewed as an MCMC analysis keeping the network topology as fixed. It enabled to explore another landscape of the parameter space. At the end, the highest likelihood values were kept among the two optimizations. In order to penalize models with too many parameters, the AIC [72] and BIC [73] criteria were adopted, using the following expressions: AIC = 2*p –* 2 ln(L) and BIC = – 2 ln(L) + *p* ln(*m*), where *m* and *p* refer to the number of sites and the number of parameters, respectively.

## Results

This section is divided in two parts: the first part is devoted to simulated data while the second part focuses on the analysis of real data.

## Simulations

In this simulation study, we compare performances of SnappNet and `MCMCBiMarkers`, regarding their ability to recover A and B networks (cf. Figure 6), already studied in [2], and the more complex C network. Last but not least, we compare the two software, in terms of CPU time and memory usage. In this case, our evaluation criterion is the likelihood computed on network C and its variants, as this step is usually repeated million times in an MCMC analysis.

### Study of networks A and B

#### 1) Ability to recover the network topology

Table 1 reports on the accuracy of SnappNet, regarding the correct topology of networks A and B. As in [2], we considered one individual for each species. Note that under this setting, population sizes *θ* corresponding to external branches are unidentifiable, as there is no coalescence event occurring along these branches. We studied different densities of markers and different priors on *θ*. Besides, we focused on either a) the true prior Γ(1, 200) with 𝔼 (*θ*) = 0.005, b) the false prior Γ(1, 1000) with 𝔼 (*θ*) = 0.001, or c) the false prior Γ(1, 2000) with 𝔼 (*θ*) = 5 *×* 10^*-*4^. Last, in order to compare our results with [2], we considered the *u* and *v* rates as known parameters in this study on topological accuracy. Indeed, `MCMCBiMarkers` relies on the operators of [55] that do not allow changes of these rates.

**Table 1.** Accuracy of SnappNet on simulated data, regarding the correct topology of networks A and B (see Figure 6). Results are given as a function of the number of sites and as a function of the hyperparameter values *α* and *β* for the prior on *θ* (*θ ∼* Γ(*α, β*) and 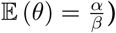. Here, one lineage was simulated per species. Constant sites are included in the analysis, the rates *u* and *v* are considered as known, and 20 replicates are considered for each simulation set up (criterion ESS*>* 200 ; *d ∼ε* (0.1), *r ∼* Beta(1, 1), *τ*_0_ *∼ε* (10) for the network prior).

| Number of sites<br>Hyperparameters | Network A |  |  | Network B |  |  |
| --- | --- | --- | --- | --- | --- | --- |
|  | 1,000 | 10,000 | 100,000 | 1,000 | 10,000 | 100,000 |
| **True** ($\alpha = 1$, $\beta = 200$, $\frac{\alpha}{\beta} = 0.005$) | 0% | 100% | 100% | 0% | 81.25% | 100% |
| **False** ($\alpha = 1$, $\beta = 1000$, $\frac{\alpha}{\beta} = 0.001$) | 0% | 94.73% | 91.30% | 0% | 80% | 95.65% |
| **False** ($\alpha = 1$, $\beta = 2000$, $\frac{\alpha}{\beta} = 5 \times 10^{-4}$) | 0% | 100% | 80% | 0% | 85% | 85.71% |

First consider simulations under the true prior. As shown in Table 1, in presence of a large number of markers, SnappNet recovered networks A and B with high accuracy. In particular, when *m* = 100, 000 sites were used, the posterior distributions were only concentrated on the true networks. For *m* = 10, 000, SnappNet’s accuracy on network B decreased slightly to 81.25%. In contrast, network A remained perfectly recovered. This is not surprising since network B is more complex than network A. Our results are similar to those of [2], who found that `MCMCBiMarkers` required 10,000 sites to infer precisely networks A and B.

However, for small number of sites (*m* = 1, 000), we observed differences between SnappNet and `MCMCBiMarkers`: SnappNet always inferred trees (see Figure 8), whereas `MCMCBiMarkers` inferred networks. For instance, on Network A, `MCMCBiMarkers` inferred the true network in approximately 75% of cases, whereas SnappNet proposed a tree resulting from removing just one branch in this network^1^, in 78.71% of the samples. Details on the trees inferred by SnappNet are given in Table 1 of the Supplementary Material.

**Fig 8.**
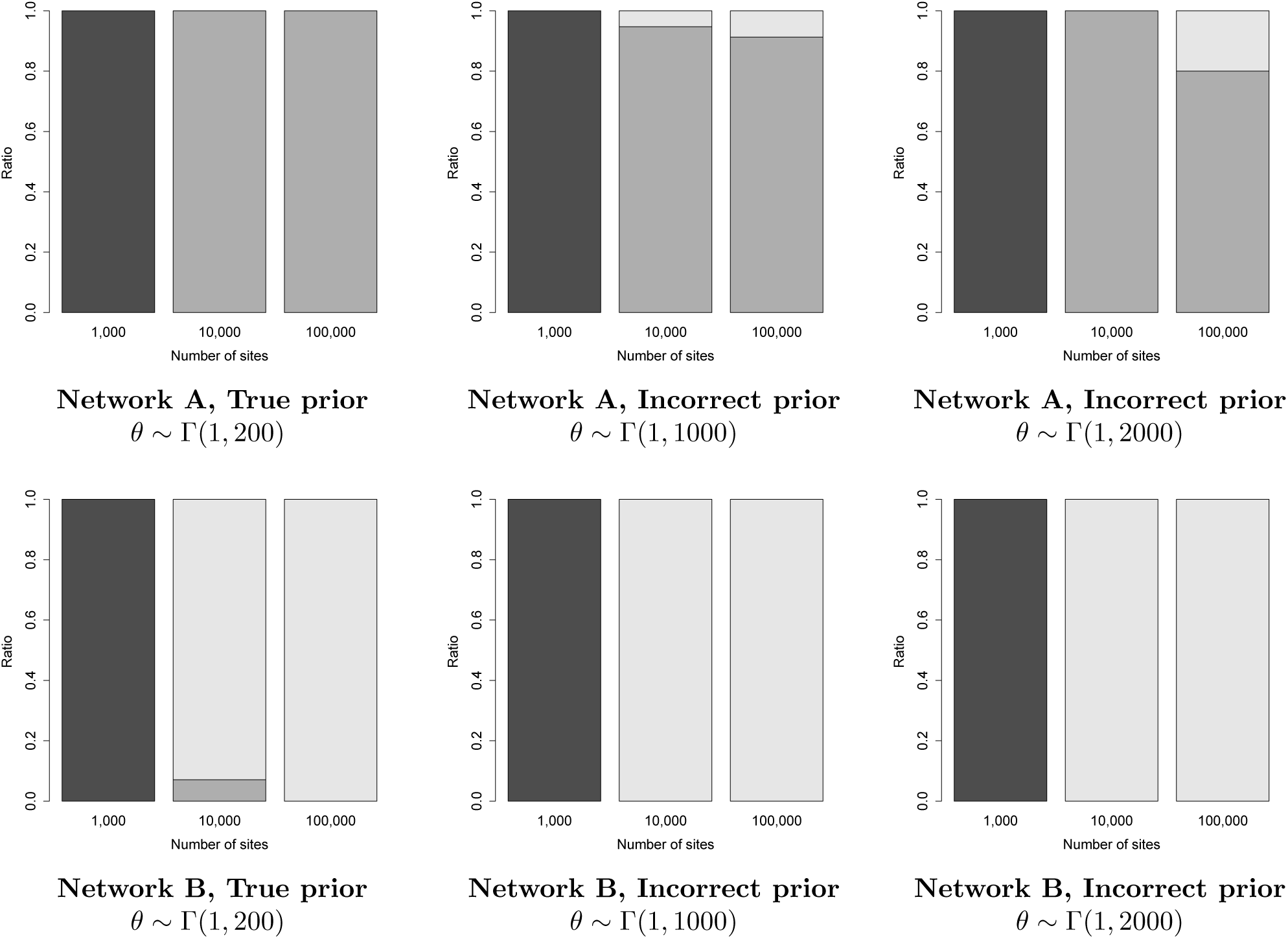
The ratio of trees (black), 1-reticulation networks (dark grey), 2-reticulations networks (light gray), sampled by SnappNet, under the different simulations settings studied in Table 1. Recall that networks A and B contain 1 and 2 reticulations, respectively.

Similarly, on network B that hosts 2 reticulations, `MCMCBiMarkers` almost always inferred a 1-reticulation network for *m* = 1, 000, whereas SnappNet hesitated mainly between two trees, trees (((Q,R),L),(A,C)) and (((Q,L),R),(A,C)), in 35.28% and 28.54% of cases, respectively. This different behavior among the two software is most likely due to the fact that their prior models differ. Anyway, with only 1,000 markers, `MCMCBiMarkers` and SnappNet were both unable to recover network B.

Now consider on simulations based on incorrect priors. This mimics real cases where there is no or little information on the network underlying the data. Recall that these priors are incorrect since 𝔼 (*θ*) is either fixed to 0.001 or 5 *×* 10^*-*4^, instead of being equal to the true value 0.005. In other words, these priors underestimate the number of ILS events. When considering as few as 1,000 sites, SnappNet only inferred trees (cf. Table 1 in Supplementary Material), whereas `MCMCBiMarkers` inferred networks. For *m* = 10, 000 and *m* = 100, 000 sites, SnappNet presented very good accuracy on network A. In the rare cases where the true network was not found, SnappNet inferred a too complex scenario with two reticulations (see Figure 8). The bias induced by incorrect priors (underestimating ILS) led the method to fit the data by adding supplementary edges to the network. Nevertheless, on network B, SnappNet’s accuracy remained fair and interestingly, for *m* = 10, 000 and *m* = 100, 000 sites, SnappNet sampled exclusively 2-reticulation networks (see Figure 8). To sum up, SnappNet’s accuracy did not really deteriorate with incorrect priors.

#### 2) Ability to estimate continuous parameters

Recall that in our modelling, the continuous parameters are branch lengths, inheritance probabilities *γ*, population sizes *θ* and instantaneous rates (*u* and *v*). As in [56], we also studied the network length and the network height, that is the sum of the branch lengths and the distance between the root node and the leaves, respectively. In order to evaluate SnappNet’s ability to estimate continuous parameters, we focused on network A (following [2]) and considered the case of two lineages in each species. Indeed, under this setting, *θ* values are now identifiable for external branches: the expected coalescent time is here *θ/*2, that is to say 2.5 *×* 10^*-*3^, which is a smaller value than all external branch lengths. In other words, a few coalescent events should happen along external branches. For these analysis, we considered exclusively the true prior on *θ* and we bounded the number of reticulations to 2 (as in [2]) in order to limit the computational burden. In the following, we consider the cases where a) input markers can be invariant or polymorphic, and b) only polymorphic sites are considered.

#### 2a) Constant sites included in the analysis

Before describing results on continuous parameters, let us first mention results regarding the topology. Although the number of lineages was increased in comparison with the previous experiment, SnappNet still sampled exclusively trees for *m* = 1, 000, and always recovered the correct topology for *m* = 10, 000 and *m* = 100, 000. Note that for *m* = 1, 000, we observed that generated data sets contained 78% invariant sites on average given the parameters of the simulation, so that such simulated data sets only contained on average 220 variable sites.

In order to limit the computational burden, the analysis for m=100,000 relied only on 17 replicates with ESS*>* 200. Figure 9 reports on the estimated network height and the estimated network length. As expected, the accuracy increased with the number of sites. Figure 10 shows the same behaviour, regarding the inheritance probability *γ*, the rates *u* and *v*. Figure 11 is complementary to Figure 9, since it reports on the estimated node heights. All node heights were estimated quite accurately, which is not surprising in view of the results on the network length. Figure 12 is dedicated to population sizes. For external branches, SnappNet’s was able to estimate *θ* values very precisely.

**Fig 9.**
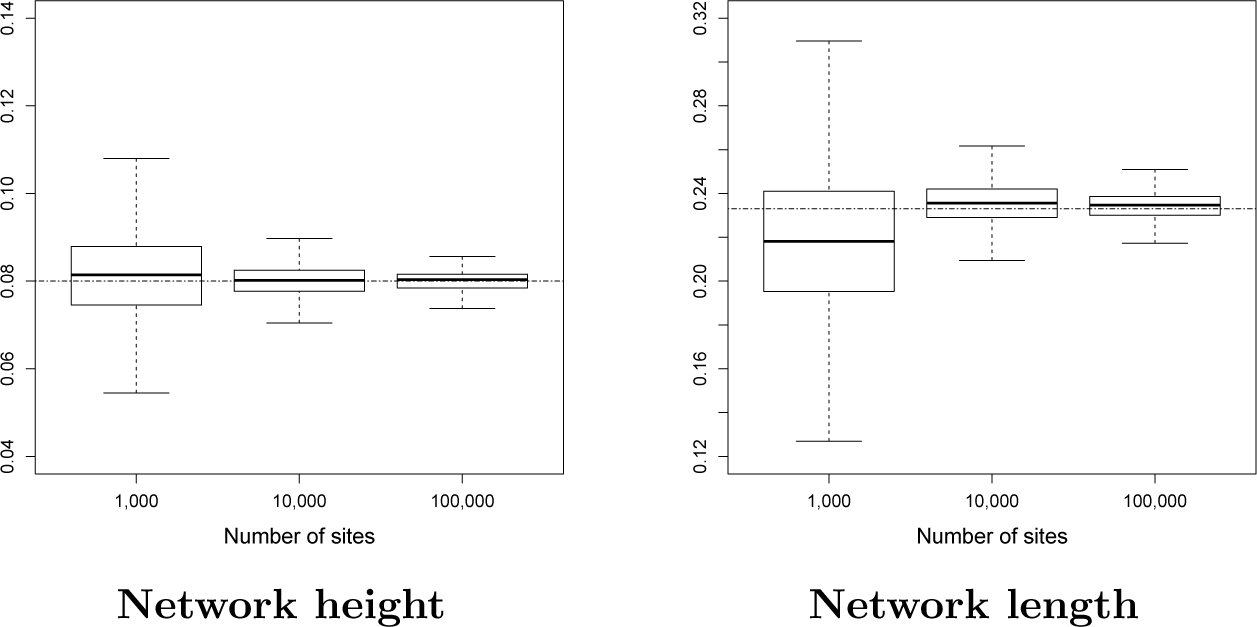
Estimated height and length for network A (see Figure 6), as a function of the number of sites. Heights and lengths are measured in units of expected number of mutations per site. True values are given by the dashed horizontal lines. Two lineages per species were simulated. Constant sites are included in the analysis, and 20 replicates are considered for each simulation set up (criterion ESS*>* 200 ; *θ ∼* Γ(1, 200), *d ∼ ε* (0.1), *r ∼* Beta(1, 1), *τ*_0_ ∼ ε (10) for the priors, number of reticulations bounded by 2 when exploring the network space).

**Fig 10.**
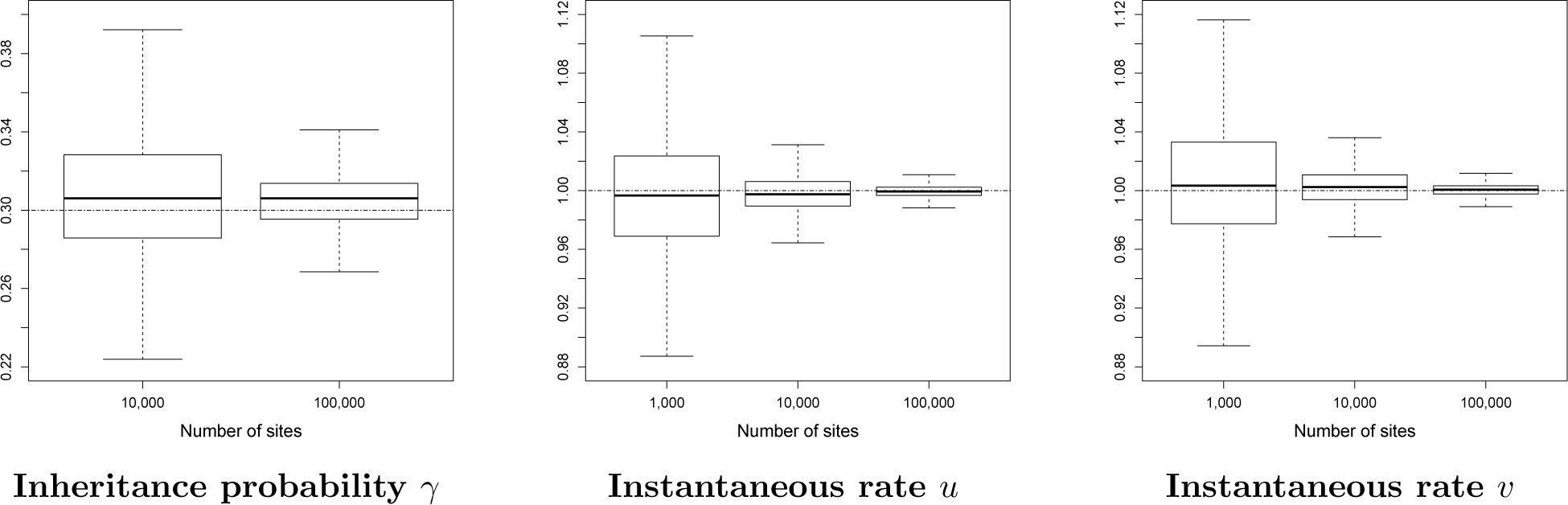
Estimated inheritance probability and instantaneous rates for network A (see Figure 6), as a function of the number of sites. True values are given by the dashed horizontal lines. Same framework as in Figure 9.

**Fig 11.**
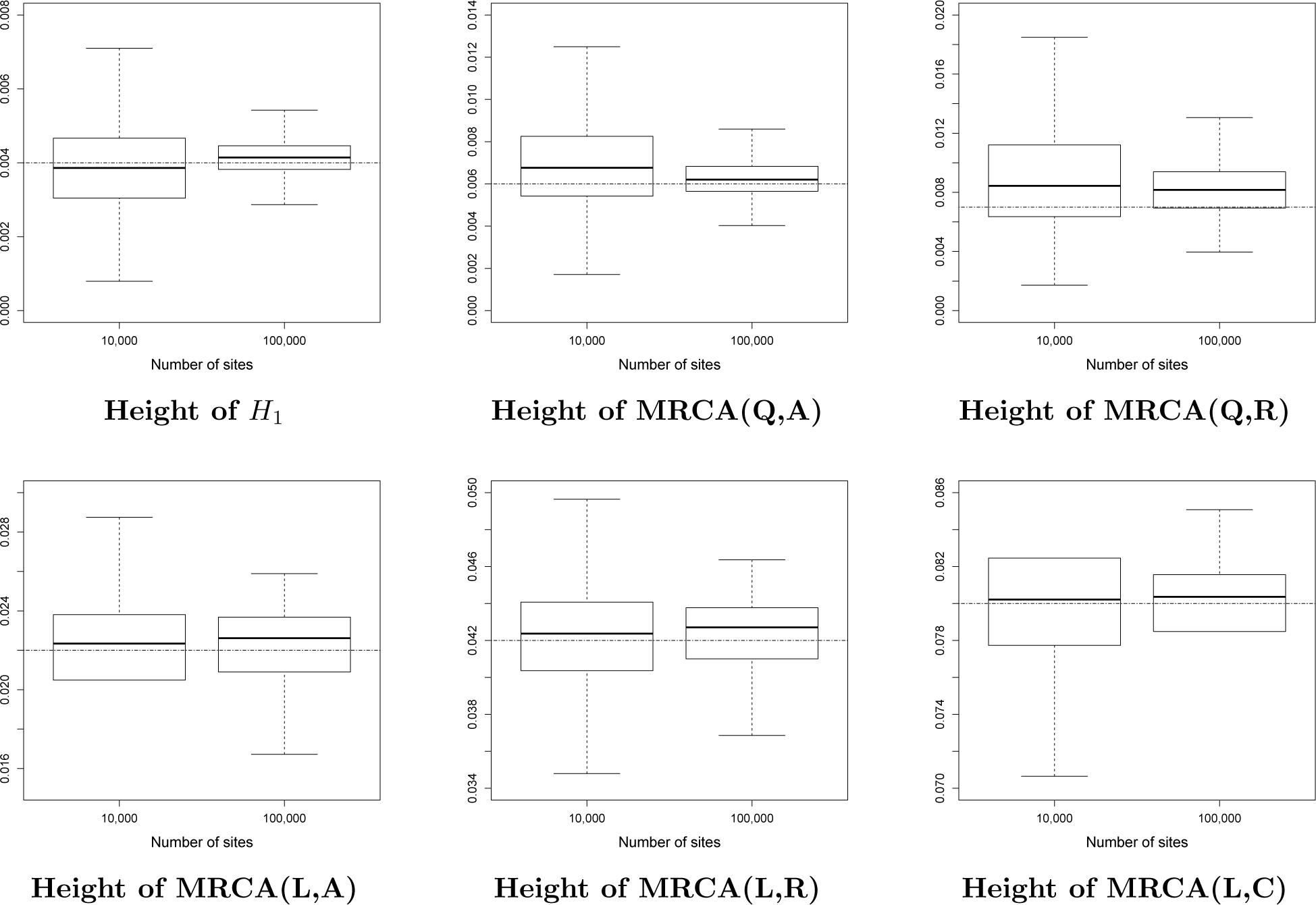
Estimated node heights of network A (see Figure 6), as a function of the number of sites. Heights are measured in units of expected number of mutations per site. True values are given by the dashed horizontal lines. Same framework as in Figure 9. The initials MRCA stand for “Most Recent Common Ancestor”.

**Fig 12.**
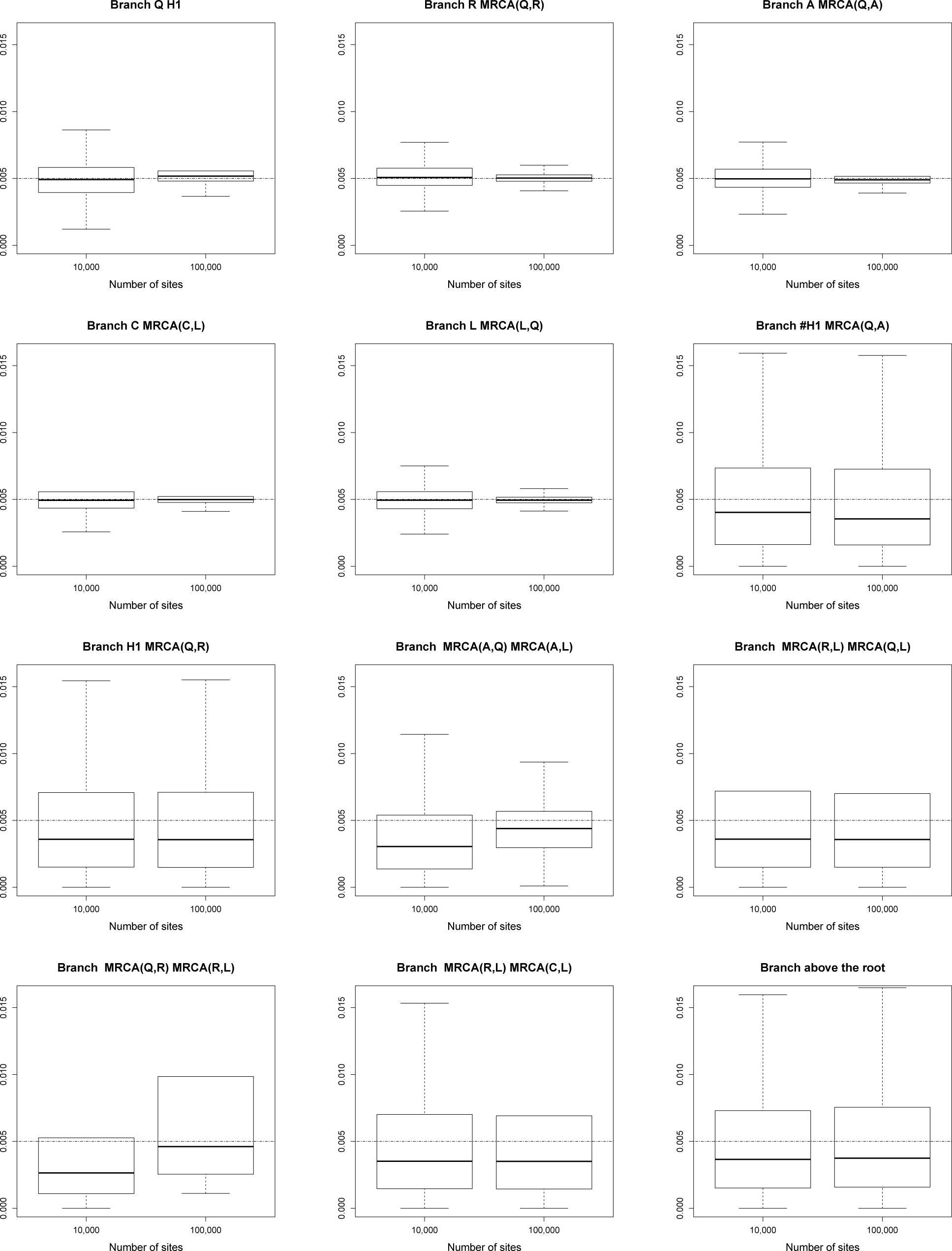
Estimated population sizes *θ* for each branch of network A (see Figure 6), as a function of the number of sites. True values are given by the dashed horizontal lines. Same framework as in Figure 9. The initials MRCA stand for “Most Recent Common Ancestor”.

Performances slightly deteriorated on internal branches (see the box plots, from number 6 to number 12) whose *θ* values were underestimated (see the medians) and showed a higher posterior variance. This phenomenon was also observed for `MCMCBiMarkers` [2, Figure 7 obtained under a different setting].

Overall, on network A, SnappNet and `MCMCBiMarkers` are two methods with comparable abilities for inferring the continuous parameters.

#### 2b) Only polymorphic sites included in the analysis

In order to control for the fact that this analysis relies only on polymorphic sites, the likelihood of the data for a network Ψ becomes a conditional likelihood equal to ℙ (*X*_1_, *…, X*_*m*_ | Ψ) */* ℙ (“the m sites are polymorphic”|Ψ), due to Bayes rules.

Before focusing on continuous parameters, let us describe results regarding the topology. As mentioned in [2], polymorphic sites help to recover the topology. For *m* = 1, 000, SnappNet now recovers the correct topology with high frequency (e.g., in 94.45% of cases). SnappNet always sampled the true network for *m* = 10, 000 and *m* = 100, 000. In order to reduce the computational burden for *m* = 100, 000, our analysis relied on the 12 replicates that achieved ESS*>* 100.

Next, the same analysis was performed without applying the correction factor ℙ(“the m sites are polymorphic” |Ψ), which is done by toggling an option within the software. For *m* = 1, 000, the accuracy dropped to 23.81%, while for *m* = 10, 000 and *m* = 100, 000, the accuracy was still high (e.g., 95.24% and 95.65%, respectively).

Again, having very few informative sites penalizes SnappNet, and using the correct likelihood computation is important here.

We also highlight that for *m* = 100, 000, the sampler efficiency (i.e. the ratio ESS/nb iterations without burnin) was much larger when the additional term was omitted (1.75 *×* 10^*-*4^ vs. 2.55 *×* 10^*-*5^). It enabled us to consider 20 replicates with ESS*>* 200 in this new experiment.

Let us move on to the estimation of continuous parameters. Figures 13, 14, 15 and 16, illustrate results obtained from the experiment incorporating the correction factor. As previously, the network height, the network length, the rates *u* and *v*, the inheritance probability *γ* and the node heights were estimated very precisely. As expected, the accuracy increased with the number of sites. Estimated *θ* values were very satisfactory for external branches, whereas a slight bias was still introduced on internal branches.

**Fig 13.**
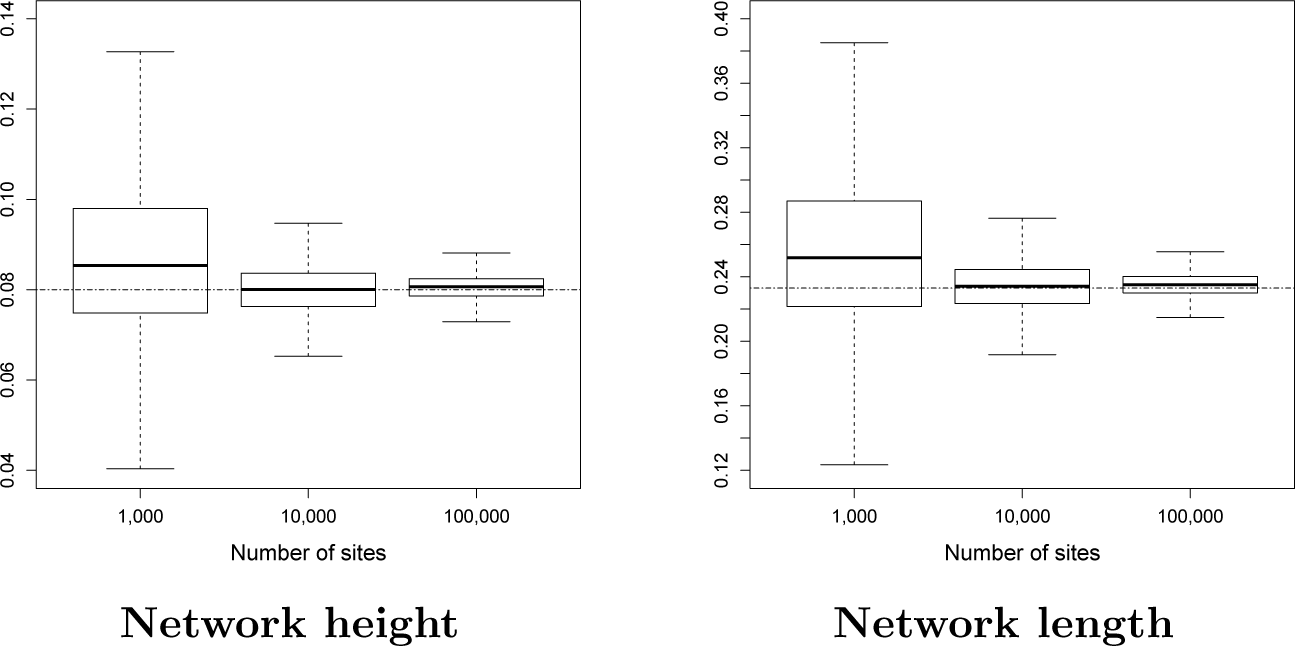
Same as Figure 9, except that only polymorphic sites are taken into account (criterion ESS*>* 200 for m=1,000 and m=10,000, and criterion ESS*>* 100 for m=100,000).

**Fig 14.**
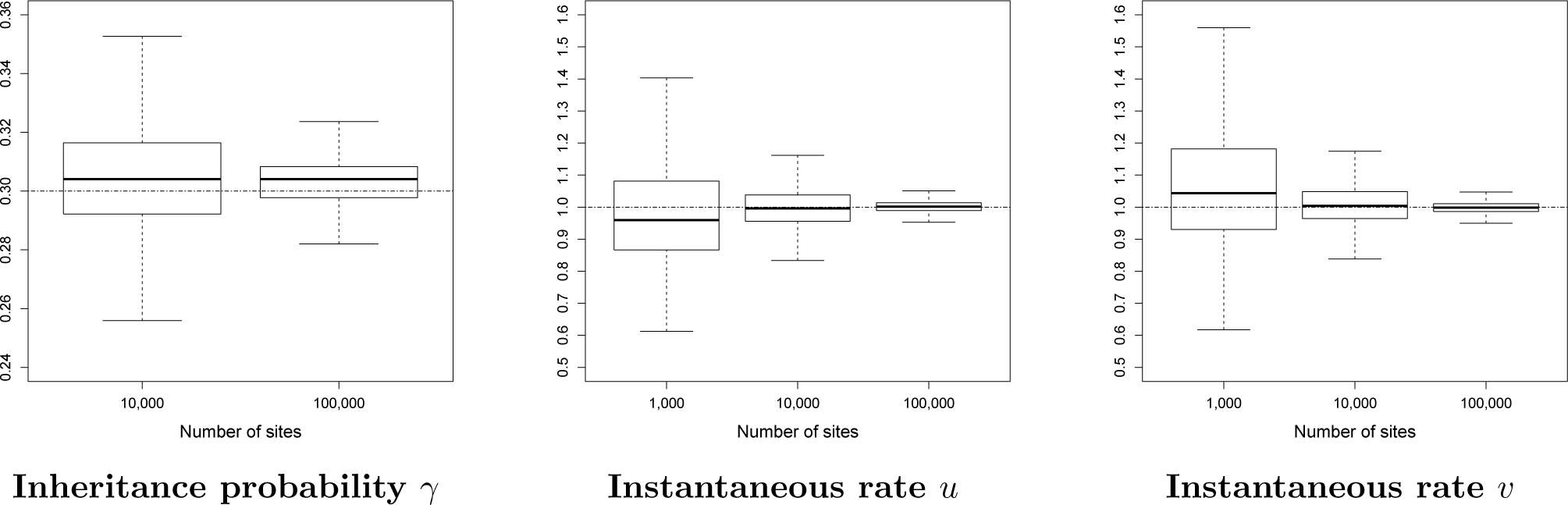
Same as Figure 10, except that only polymorphic sites are taken into account.

**Fig 15.**
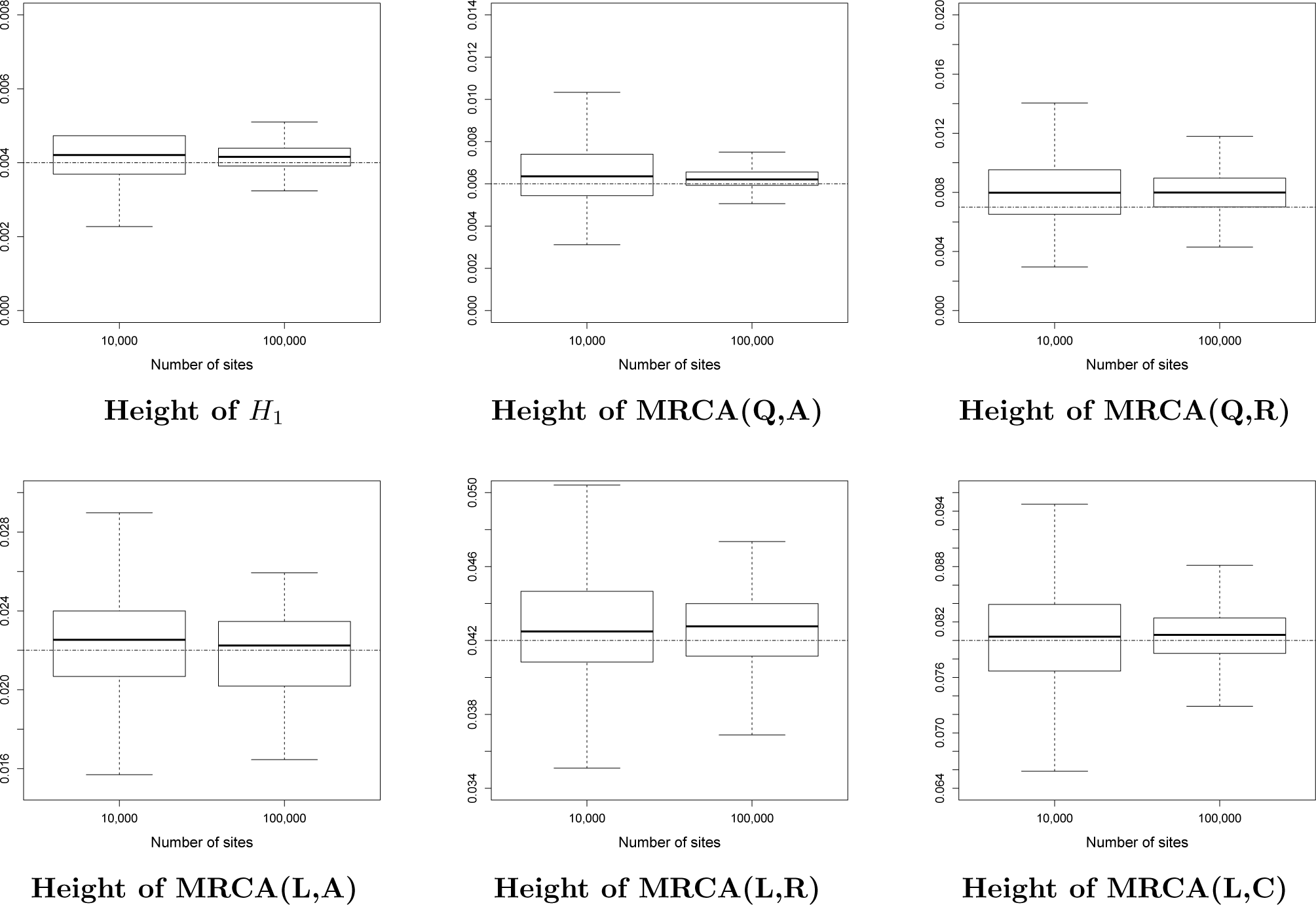
Same as Figure 11, except that only polymorphic sites are taken into account.

**Fig 16.**
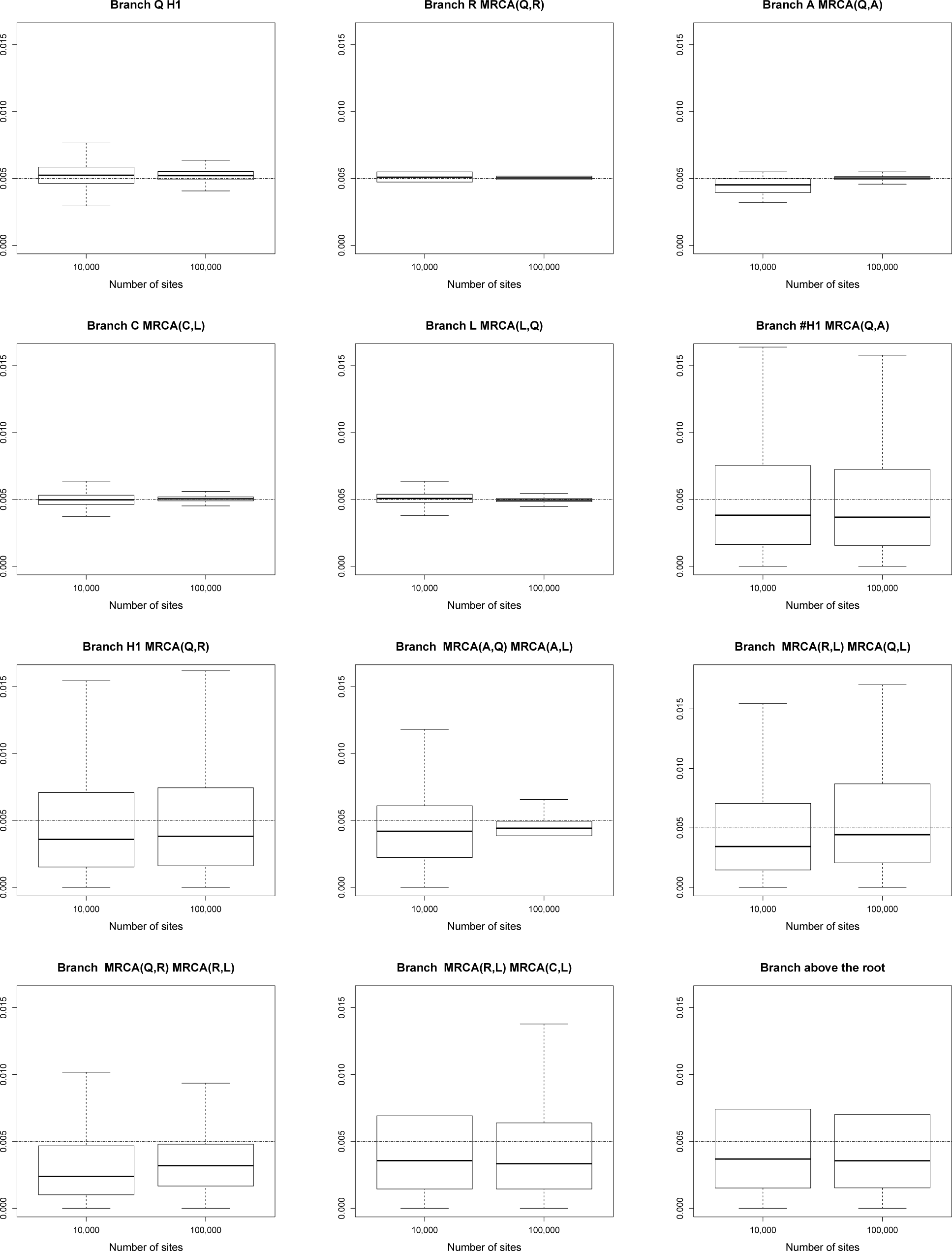
Same as Figure 12, except that only polymorphic sites are taken into account.

Last, for the analysis without the correction factor, we observed a huge bias regarding network height and network length (cf Figure 4 in Supplementary Material). Surprisingly, the rates *u* and *v* were still very accurately estimated.

### Study of network C and its variants

We focus here on network C and its variants, that is networks containing reticulation nodes on top of one another.

#### 1) Ability to recover the network topology

Tables 2 and 3 report respectively SnappNet and `MCMCBiMarkers` performances, regarding their ability to recover the correct topology of network C. Recall that network C is a level 2 network. We considered one lineage in species O, A and D, and we let the number of lineages in species C and D vary. We studied either a) 1 lineage, or b) 4 lineages, in these hybrid species. In order to limit the computational burden for SnappNet, the ESS criterion was decreased to 100 and the number of reticulations was also bounded by 2.

**Table 2.**
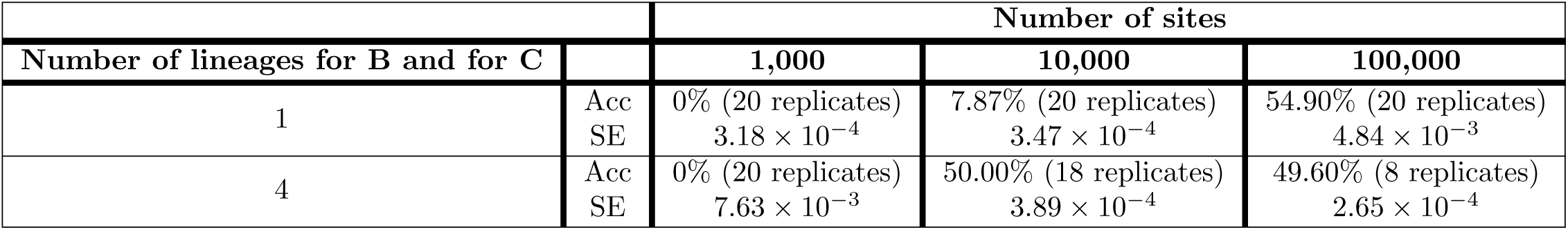
Accuracy (Acc) of SnappNet on simulated data, regarding the correct topology of network C (see Figure 6). Results are given as a function of the number of sites and as a function of the number of sampled lineages in hybrid species B and C. Only one lineage was sampled in each other species. Constant sites are included in the analysis and the rates *u* and *v* are considered as known. Accuracy is computed on the basis of replicates for which the criterion ESS*>* 100 is fulfilled. The sampler efficiency (SE) is also indicated (true hyperparameter values for the prior on *θ*, i.e. *θ ∼* Γ(1, 200) ; as a network prior *d ∼ε* (0.1), *r ∼* Beta(1, 1), *τ*_0_ *∼ε* (10) ; number of reticulations bounded by 2 when exploring the network space).

|  |  | Number of sites |  |  |
| --- | --- | --- | --- | --- |
| Number of lineages for B and for C |  | 1,000 | 10,000 | 100,000 |
| 1 | Acc | 0% (20 replicates) | 7.87% (20 replicates) | 54.90% (20 replicates) |
| | SE | $3.18 \times 10^{-4}$ | $3.47 \times 10^{-4}$ | $4.84 \times 10^{-3}$ |
| 4 | Acc | 0% (20 replicates) | 50.00% (18 replicates) | 49.60% (8 replicates) |
| | SE | $7.63 \times 10^{-3}$ | $3.89 \times 10^{-4}$ | $2.65 \times 10^{-4}$ |

**Table 3.** Accuracy (Acc) of MCMCBiMarkers on simulated data, regarding the correct topology of network C (see Figure 6). As in Table 2, results are given as a function of the number of sites and as a function of the number of sampled lineages in hybrid species B and C. 1 lineage is considered in other species, constant sites are included in the analysis, and the rates *u* and *v* are considered as known. 1.5 *×* 10^6^ iterations are considered. 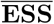 is the average ESS over the different replicates, and SE stands for the sampler efficiency.

| Number of lineages for B and for C |  | Number of sites |  |  |
| --- | --- | --- | --- | --- |
|  |  | 1,000 | 10,000 | 100,000 |
| 1 | Acc | 0% (20 replicates) | 4.84% (20 replicates) | 0% (20 replicates) |
| | SE | $9.70 \times 10^{-5}$ | $3.10 \times 10^{-5}$ | $3.60 \times 10^{-5}$ |
| | $\overline{\text{ESS}}$ | 126.08 | 40.38 | 46.80 |
| 4 | Acc | 0% (20 replicates) | 0% (12 replicates) | 0% (9 replicates) |
| | SE | $2.38 \times 10^{-4}$ | $8.53 \times 10^{-5}$ | $1.03 \times 10^{-5}$ |
| | $\overline{\text{ESS}}$ | 309.00 | 110.90 | 159.96 |

In order to mimic exactly what has been done in [2], we let `MCMCBiMarkers` run during 1,500,000 interations instead of adopting an ESS criterion. Data were simulated thanks to `simBiMarker` [2]. Indeed, as its cousin SimSnapp, SimSnappNet generates only count data (the number of alleles per site and per species). In contrast, `simBiMarker` generates actual sequences, a prerequisite for running `MCMCBiMarkers`. The commands used under the 4 lineages scenario, are described in Section 4 of the Supplementary Material. Note that for calibrating `MCMCBiMarkers` network prior, the maximum number of reticulations was set to 2, and the prior Poisson distribution on the number of reticulation nodes was centered on 2.

According to Table 2, as expected, SnappNet’s accuracy increased with the number of sites and with the number of lineages in the hybrid species. For instance, in presence of one lineage in hybrid species B and C, the accuracy increased from 7.87% for m=10,000 to 54.90% for m=100,000. In the same way, when 4 lineages were considered instead of a unique lineage, we observed an increase from 7.87% to 50.00% for m=10,000. Note that the accuracy of 49.60% reported for m=100,000 and 4 lineages, is based only on 8 replicates. This accuracy should have increased if we had let SnappNet run longer to obtain results from more replicates.

Surprisingly, in most cases studied, `MCMCBiMarkers` was unable to recover the true topology of network C. The different behaviors of `MCMCBiMarkers` and SnappNet, must be due to the fact that `MCMCBiMarkers` incorporates a network prior similar to ancestral recombination graphs, whereas SnappNet considers the birth hybridization process of [56]. Nevertheless, the ratios of trees, 1-reticulation networks and 2-reticulations networks, sampled by the two methods, were globally similar (cf. Figure 17). We first suspected that `MCMCBiMarkers` lack of performance was due to small ESS values, specially when only one lineage was sampled in hybrid species B and C. However, when the number of iterations was increased from 1.5 *×* 10^6^ to 12 *×* 10^6^, `MCMCBiMarkers` was still unable to recover network C, despite larger ESS values (see Table 2 in Supplementary Material). As a consequence, the birth hybridization process prior [56] on which SnappNet relies seems to be the best choice for recovering network C.

**Fig 17.**
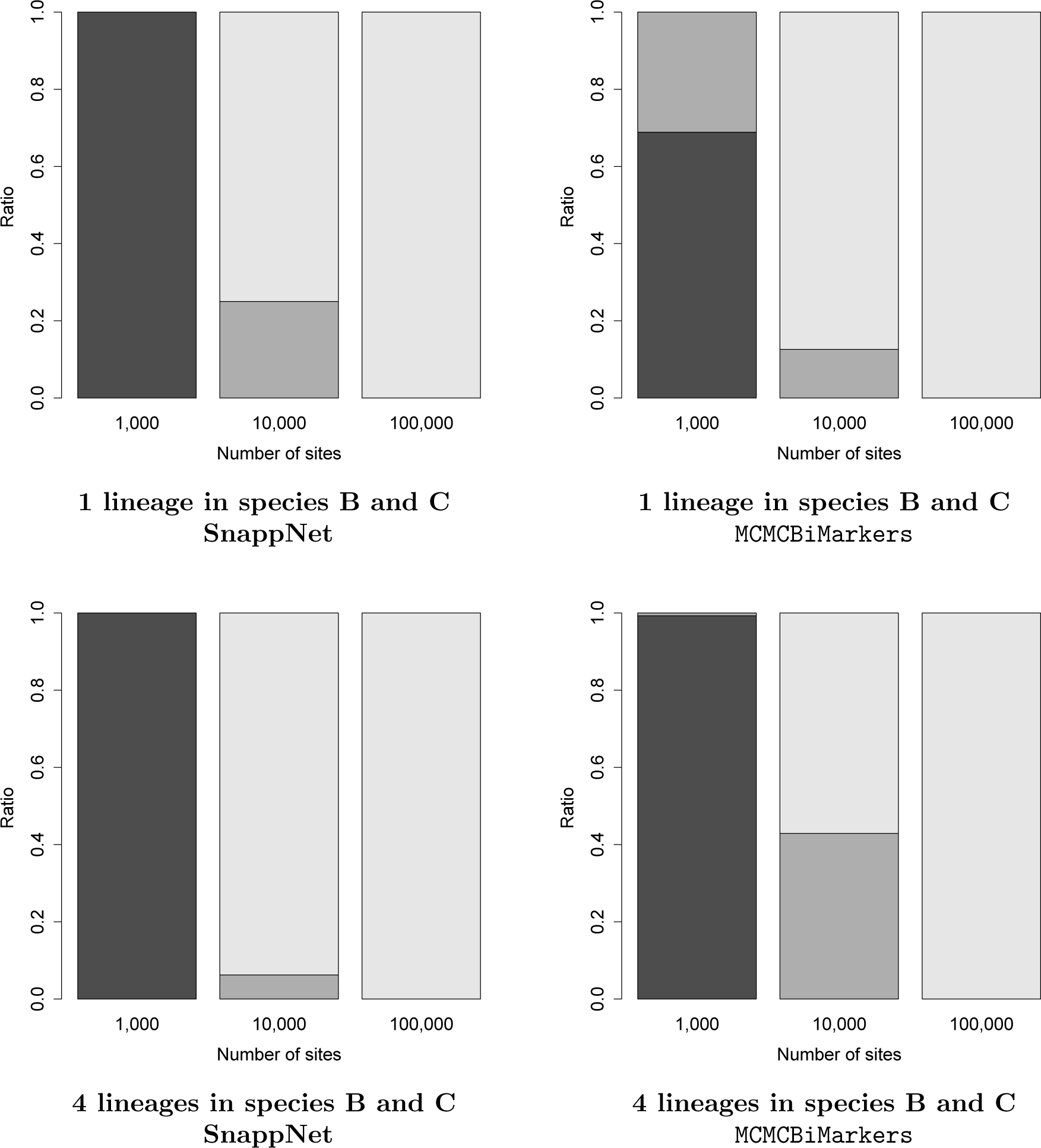
The ratio of trees (black), 1-reticulation networks (dark grey), 2-reticulations networks (light gray), sampled by SnappNet and `MCMCBiMarkers`, when data were simulated from Network C (see Tables 2 and 3). Recall that network C contains 2 reticulations.

#### 2) CPU time and required memory

To compare the CPU time and memory required by SnappNet and `MCMCBiMarkers` on a single likelihood calculation, we focused on network C (see Figure 6), with and without outgroup (i.e. the species O), and networks C(3) and C(4), again with and without outgroup (see Figure 7). The simulations protocol used here is similar to that used in the previous sections, where here we fixed 10 lineages in species C and one lineage in the other species, *m* = 1, 000 sites and 20 replicates per each network. The likelihood calculations were run on the true network.

The experiments were executed on a full quad socket machine with a total of 512GB of RAM (4 * 2.3 GHz AMD Opteron 6376 with 16 Cores, each with a RDIMM 32Go Quad Rank LV 1333MHz processor). The jobs that did not finished within two weeks, or required more than 128 GB, were discarded.

The results are reported in Tables 4, 5 and 6. From the tables, we can see that SnappNet managed to run for all the scenarios within the two weeks limit: on average within 2.6 minutes and using 1.66 GB on network C without outgroup, within 5.6 minutes and using 2 GB on network C with outgroup, within 14.2 minutes and using 2.2 GB on network C(3) without outgroup, within 24.7 minutes and using 2.2 GB on network C(3) with outgroup, within 45.46 minutes and using 2.6 GB on network C(4) without outgroup, and finally, within 70.9 minutes and using 3.1 GB on network C(4) with outgroup.

**Table 4.** Comparison regarding CPU time and memory, between SnappNet and MCMCBiMarkers. The evaluation criterion is the likelihood calculation. The focus is on network C and 10 lineages are present in species C and 1 lineage in other species. Each row corresponds to a different replicate.

|  | CPU time |  | Memory |  |
| --- | --- | --- | --- | --- |
|  | SnappNet<br>(in minutes) | MCMCbiMarkers<br>(in hours) | SnappNet<br>(max in GB) | MCMCbiMarkers<br>(max in GB) |
| Network C Without Outgroup O | 2.63 | 15.51 | 1.66 | 8.75 |
|  | 2.58 | 14.52 | 1.66 | 8.78 |
|  | 2.52 | 13.79 | 1.66 | 8.75 |
|  | 2.61 | 15.21 | 1.66 | 8.80 |
|  | 2.65 | 14.47 | 1.65 | 8.79 |
|  | 2.59 | 13.90 | 1.66 | 8.77 |
|  | 2.62 | 14.44 | 1.68 | 8.77 |
|  | 2.60 | 14.35 | 1.66 | 8.77 |
|  | 2.65 | 14.97 | 1.66 | 8.77 |
|  | 2.63 | 14.00 | 1.66 | 8.76 |
|  | 2.68 | 14.61 | 1.66 | 8.74 |
|  | 2.63 | 15.01 | 1.67 | 8.78 |
|  | 2.62 | 14.45 | 1.80 | 8.75 |
|  | 2.64 | 13.91 | 1.66 | 8.76 |
|  | 2.59 | 14.27 | 1.65 | 8.76 |
|  | 2.66 | 15.14 | 1.65 | 8.75 |
|  | 2.62 | 14.53 | 1.64 | 8.76 |
|  | 2.70 | 15.42 | 1.67 | 8.75 |
|  | 2.58 | 14.51 | 1.65 | 8.75 |
|  | 2.65 | 14.53 | 1.65 | 8.75 |
| Network C | 5.56 | 35.94 | 1.96 | 8.79 |
|  | 5.68 | 34.24 | 1.96 | 8.78 |
|  | 5.73 | 32.65 | 1.96 | 8.82 |
|  | 5.45 | 34.20 | 1.96 | 8.78 |
|  | 5.60 | 33.24 | 1.96 | 8.79 |
|  | 5.58 | 31.99 | 1.96 | 8.77 |
|  | 5.36 | 32.41 | 1.97 | 8.78 |
|  | 5.48 | 34.40 | 1.95 | 8.78 |
|  | 5.91 | 31.12 | 1.97 | 8.82 |
|  | 5.98 | 34.36 | 1.98 | 8.80 |
|  | 5.54 | 33.60 | 2.19 | 8.79 |
|  | 5.45 | 33.81 | 1.95 | 8.80 |
|  | 5.69 | 31.89 | 1.96 | 8.77 |
|  | 5.46 | 31.42 | 1.95 | 8.77 |
|  | 5.68 | 34.54 | 1.95 | 8.83 |
|  | 5.55 | 35.24 | 2.21 | 8.80 |
|  | 5.82 | 33.44 | 1.97 | 8.77 |
|  | 5.64 | 32.49 | 1.97 | 8.82 |
|  | 5.72 | 34.87 | 2.21 | 8.81 |
|  | 5.71 | 33.30 | 1.96 | 8.82 |

**Table 5.** Comparison regarding CPU time and memory, between SnappNet and MCMCBiMarkers. The evaluation criterion is the likelihood calculation. The focus is on network C(3) (see Figure 7) and 10 lineages are present in species C1 and C2, and 1 lineage in other species. Each row corresponds to a different replicate.

|  | CPU time |  | Memory |  |
| --- | --- | --- | --- | --- |
|  | SnappNet<br>(in minutes) | MCMC*BiMarkers*<br>(in hours) | SnappNet<br>(max in GB) | MCMC*BiMarkers*<br>(max in GB) |
| Network C(3) Without Outgroup | 13.57 | > 336 | 2.19 | < 64 |
|  | 14.98 |  | 2.20 |  |
|  | 14.08 |  | 2.19 |  |
|  | 13.87 |  | 2.19 |  |
|  | 14.20 |  | 2.21 |  |
|  | 14.66 |  | 2.18 |  |
|  | 14.77 |  | 2.19 |  |
|  | 13.40 |  | 2.18 |  |
|  | 14.60 |  | 2.18 |  |
|  | 14.29 |  | 2.20 |  |
|  | 13.44 |  | 2.18 |  |
|  | 13.31 |  | 2.19 |  |
|  | 14.18 |  | 2.19 |  |
|  | 14.94 |  | 2.18 |  |
|  | 14.17 |  | 2.18 |  |
|  | 14.91 |  | 2.20 |  |
|  | 14.38 |  | 2.18 |  |
|  | 13.37 |  | 2.18 |  |
|  | 14.80 |  | 2.17 |  |
|  | 14.27 |  | 2.18 |  |
| Network C(3) With Outgroup | 23.75 | > 336 | 2.19 | < 64 |
|  | 25.14 |  | 2.19 |  |
|  | 25.31 |  | 2.20 |  |
|  | 24.90 |  | 2.19 |  |
|  | 25.15 |  | 2.19 |  |
|  | 24.72 |  | 2.44 |  |
|  | 25.49 |  | 2.19 |  |
|  | 24.58 |  | 2.20 |  |
|  | 23.15 |  | 2.20 |  |
|  | 25.04 |  | 2.20 |  |
|  | 24.85 |  | 2.19 |  |
|  | 24.27 |  | 2.19 |  |
|  | 24.60 |  | 2.21 |  |
|  | 24.28 |  | 2.19 |  |
|  | 25.16 |  | 2.19 |  |
|  | 23.47 |  | 2.20 |  |
|  | 24.60 |  | 2.21 |  |
|  | 24.66 |  | 2.19 |  |
|  | 25.32 |  | 2.21 |  |
|  | 25.39 |  | 2.20 |  |

**Table 6.**
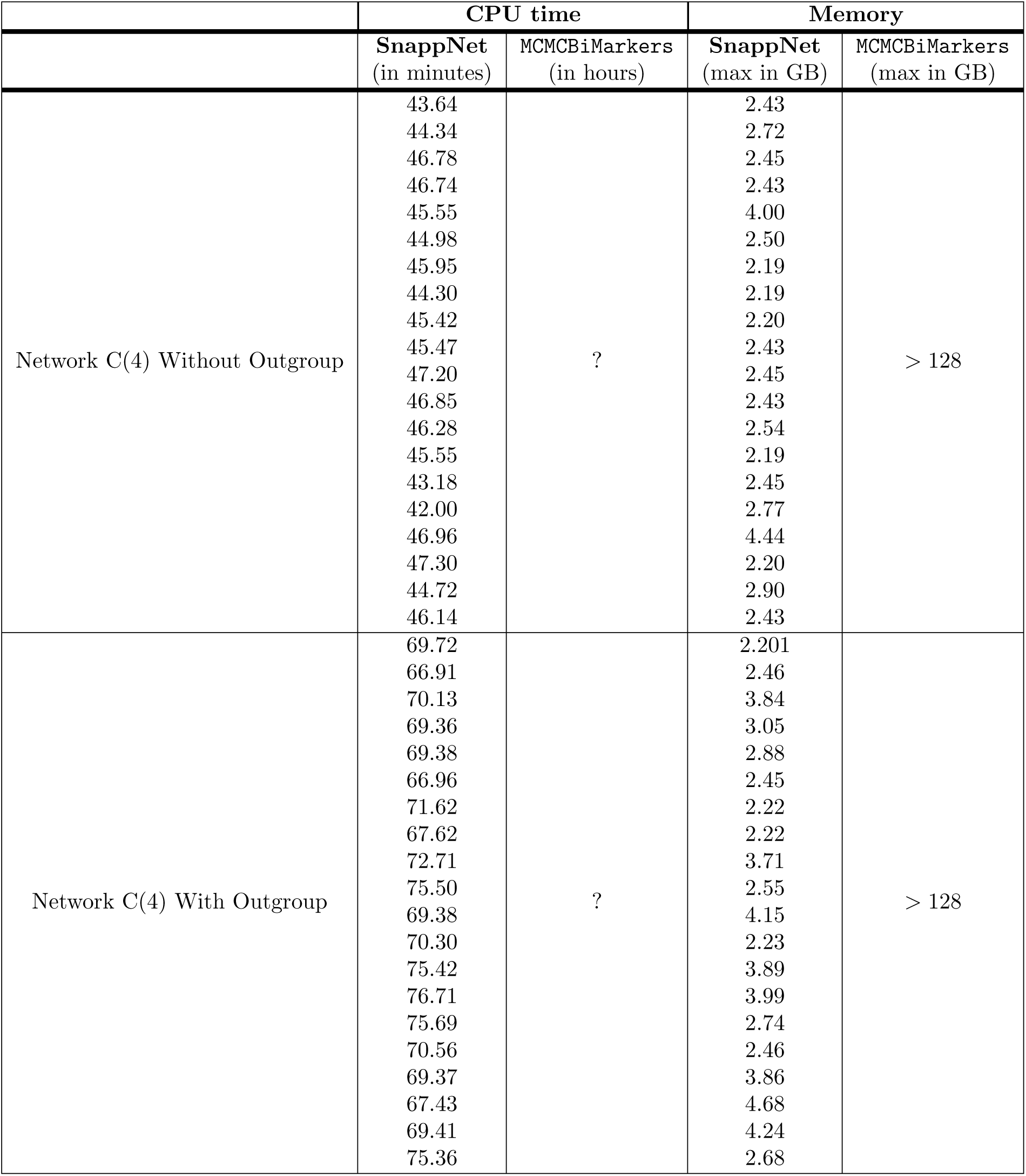
Comparison regarding CPU time and memory, between SnappNet and MCMCBiMarkers. The evaluation criterion is the likelihood calculation. The focus is on network C(4) (see Figure 7) and 10 lineages are present in species C1 C2 and C3, and 1 lineage in other species. Each row corresponds to a different replicate.

|  | CPU time |  | Memory |  |
| --- | --- | --- | --- | --- |
|  | SnappNet<br>(in minutes) | MCMC*BiMarkers*<br>(in hours) | SnappNet<br>(max in GB) | MCMC*BiMarkers*<br>(max in GB) |
| Network C(4) Without Outgroup | 43.64 | ? | 2.43 | > 128 |
|  | 44.34 |  | 2.72 |  |
|  | 46.78 |  | 2.45 |  |
|  | 46.74 |  | 2.43 |  |
|  | 45.55 |  | 4.00 |  |
|  | 44.98 |  | 2.50 |  |
|  | 45.95 |  | 2.19 |  |
|  | 44.30 |  | 2.19 |  |
|  | 45.42 |  | 2.20 |  |
|  | 45.47 |  | 2.43 |  |
|  | 47.20 |  | 2.45 |  |
|  | 46.85 |  | 2.43 |  |
|  | 46.28 |  | 2.54 |  |
|  | 45.55 |  | 2.19 |  |
|  | 43.18 |  | 2.45 |  |
|  | 42.00 |  | 2.77 |  |
|  | 46.96 |  | 4.44 |  |
|  | 47.30 |  | 2.20 |  |
|  | 44.72 |  | 2.90 |  |
|  | 46.14 |  | 2.43 |  |
| Network C(4) With Outgroup | 69.72 | ? | 2.201 | > 128 |
|  | 66.91 |  | 2.46 |  |
|  | 70.13 |  | 3.84 |  |
|  | 69.36 |  | 3.05 |  |
|  | 69.38 |  | 2.88 |  |
|  | 66.96 |  | 2.45 |  |
|  | 71.62 |  | 2.22 |  |
|  | 67.62 |  | 2.22 |  |
|  | 72.71 |  | 3.71 |  |
|  | 75.50 |  | 2.55 |  |
|  | 69.38 |  | 4.15 |  |
|  | 70.30 |  | 2.23 |  |
|  | 75.42 |  | 3.89 |  |
|  | 76.71 |  | 3.99 |  |
|  | 75.69 |  | 2.74 |  |
|  | 70.56 |  | 2.46 |  |
|  | 69.37 |  | 3.86 |  |
|  | 67.43 |  | 4.68 |  |
|  | 69.41 |  | 4.24 |  |
|  | 75.36 |  | 2.68 |  |

We were able to run `MCMCBiMarkers` for all replicates of the network C, and we can thus compare its performance with that of SnappNet. From Table 4, we see that SnappNet is remarkably faster that `MCMCBiMarkers`, needing on average only 0.29% of the time and 21% of the memory required by `MCMCBiMarkers`. `MCMCBiMarkers` needed more than 2 weeks for all scenarios on the C(3) network (requiring less than 64 GB), thus no run time is available for these scenarios. The same holds for the C(4) network scenarios, but for a different reason: all runs needed more than 128 GB each, and were discarded.

### Real data analysis

To conclude this study, we propose to illustrate SnappNet on rice real data. Diversity among Asian rice cultivars is structured around two major types which display worldwide distributions, namely Japonica and Indica, and two types localised around the Himalayas, namely *circum* Aus (cAus) and *circum* Basmati (cBasmati) [71, 74]. Japonica and Indica each have several subgroups with geographical contrast (see [71] as the most detailed description). Domestication scenarios that have been put forwards since the availability of whole genome sequences propose one to three domestications corresponding either to an early pivotal process in Japonica [75], or to multiple parallel dynamics in Japonica, Indica and cAus [14, 30], depending on whether they consider the contribution of domestication alleles by the Japonica origin as predominant or as one among others. cBasmati has been posited as a specific lineage within Japonica [75] or as a secondary derivative from admixture between Japonica and a local wild rice close to cAus [76], or between Japonica and cAus with the contribution of one or several additional cryptic sources [77].

We selected rice accessions that span all the components of global classifications and have deep genome sequencing data, and made two data sets of similar representative constitution, including three Japonica, three Indica, two cAus and three cBasmati accessions (see the *Material and methods* section). Furthermore, we considered two samplings of 12k SNPs along the whole genome alignment.

Unfortunately, our Bayesian analyses with a free exploration of the topology space led to poor mixing throughout the support of the posterior density: the chains were stuck in certain regions of the state space (see for instance [78, 79]). To overcome this issue, we focused exclusively on the 10 network topologies described in Figure 18, that we fixed in separate analyses. We computed with SnappNet the likelihoods of the different networks, and we penalized more complex models with to the AIC [72] and BIC [73] criteria.

**Fig 18.**
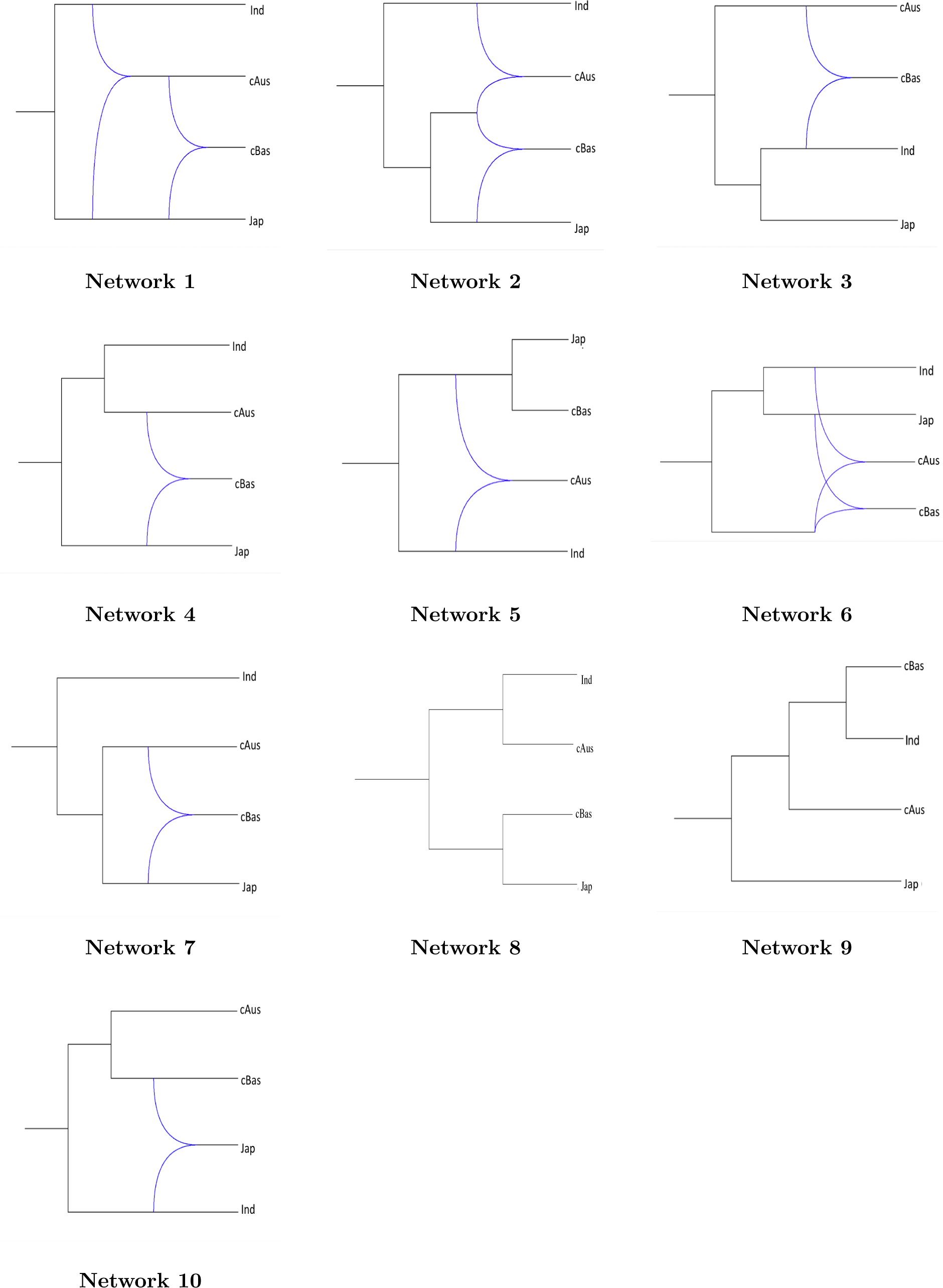
The 10 studied phylogenetic networks. Each network illustrates a different rice evolution scenario.

Tables 7 and 8, associated to the two accession samplings, report the log likelihoods, the AIC and the BIC values, for the different studied networks. For each criterion and for each data set, the ranking between networks is presented in brackets. The average ranking (cf. Table 4 in Supplementary Material), based on all these rankings, is the following: Net 1 *>* Net 7 *>* Net 2 *>* Net 4 *>* Net 10 *>* Net 5 *>* Net 6 *>* Net 8 *>* Net 3 *>* Net 9.

**Table 7.** Log Likelihood, AIC and BIC evaluations for the 10 different networks of Figure ??, each one illustrating a rice evolution process. The network ranking, given into brackets, is obtained by minimizing either the AIC or the BIC criterion. The two data sets consist in m=12,000 SNPs spread out along the rice genome. 11 rice varieties were selected for each data set.

| Network | Nb Parameters | Data set 1 |  |  | Data set 2 |  |  |
| --- | --- | --- | --- | --- | --- | --- | --- |
|  |  | Log Likelihood | AIC | BIC | Log Likelihood | AIC | BIC |
| 1 | 29 | -41699.09 | 83456.18 (2) | 83670.57 (3) | -38860.03 | 77778.07 (1) | 77992.45 (3) |
| 7 | 22 | -41724.92 | 83493.84 (3) | 83656.48 (2) | -38878.57 | 77801.14 (4) | 77963.78 (2) |
| 4 | 22 | -41669.73 | 83383.46 (1) | 83546.10 (1) | -39036.92 | 78117.84 (6) | 78280.48 (6) |
| 2 | 29 | -41755.40 | 83568.81 (4) | 83783.19 (4) | -38867.12 | 77792.24 (2) | 78006.63 (4) |
| 10 | 22 | -41936.12 | 83916.24 (7) | 84078.88 (7) | -38878.48 | 77800.96 (3) | 77963.60 (1) |
| 5 | 22 | -41921.53 | 83887.06 (6) | 84049.70 (6) | -38964.73 | 77973.46 (5) | 78136.10 (5) |
| 6 | 29 | -41874.35 | 83806.70 (5) | 84021.09 (5) | -39040.20 | 78138.40 (7) | 78352.79 (7) |
| 3 | 22 | -44264.72 | 88573.44 (8) | 88736.08 (8) | -41197.22 | 82438.44 (9) | 82601.08 (9) |
| 8 | 15 | -45586.44 | 91202.87 (10) | 91313.76 (10) | -39209.63 | 78449.26 (8) | 78560.15 (8) |
| 9 | 15 | -45041.92 | 90113.84 (9) | 90224.73 (9) | -41498.45 | 83026.9 (10) | 83137.79 (10) |

**Table 8.**
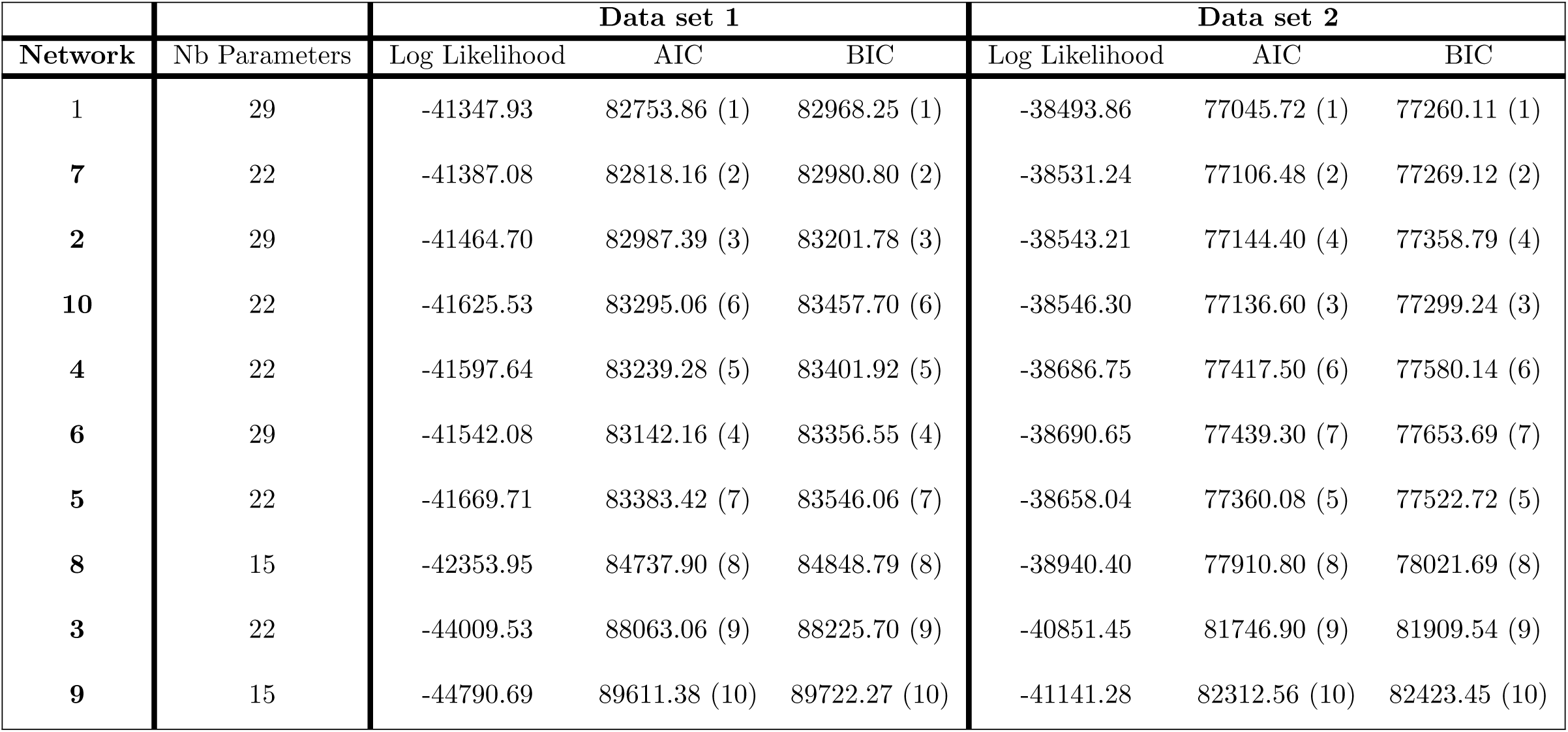
Same as Table 7 except that the analysis relies on m=12, 000 other SNPs sampled along the genome. The 11 selected rice varieties remain unchanged for the two data sets.

| Network | Nb Parameters | Data set 1 |  |  | Data set 2 |  |  |
| --- | --- | --- | --- | --- | --- | --- | --- |
|  |  | Log Likelihood | AIC | BIC | Log Likelihood | AIC | BIC |
| 1 | 29 | -41347.93 | 82753.86 (1) | 82968.25 (1) | -38493.86 | 77045.72 (1) | 77260.11 (1) |
| 7 | 22 | -41387.08 | 82818.16 (2) | 82980.80 (2) | -38531.24 | 77106.48 (2) | 77269.12 (2) |
| 2 | 29 | -41464.70 | 82987.39 (3) | 83201.78 (3) | -38543.21 | 77144.40 (4) | 77358.79 (4) |
| 10 | 22 | -41625.53 | 83295.06 (6) | 83457.70 (6) | -38546.30 | 77136.60 (3) | 77299.24 (3) |
| 4 | 22 | -41597.64 | 83239.28 (5) | 83401.92 (5) | -38686.75 | 77417.50 (6) | 77580.14 (6) |
| 6 | 29 | -41542.08 | 83142.16 (4) | 83356.55 (4) | -38690.65 | 77439.30 (7) | 77653.69 (7) |
| 5 | 22 | -41669.71 | 83383.42 (7) | 83546.06 (7) | -38658.04 | 77360.08 (5) | 77522.72 (5) |
| 8 | 15 | -42353.95 | 84737.90 (8) | 84848.79 (8) | -38940.40 | 77910.80 (8) | 78021.69 (8) |
| 3 | 22 | -44009.53 | 88063.06 (9) | 88225.70 (9) | -40851.45 | 81746.90 (9) | 81909.54 (9) |
| 9 | 15 | -44790.69 | 89611.38 (10) | 89722.27 (10) | -41141.28 | 82312.56 (10) | 82423.45 (10) |

The network that represents the simple phenetic tree derived from global molecular distances (Network 8) ranks only in the eighth position, confirming the importance of genetic exchanges in rice. The most likely scenarios 1) place the split between ancestors of Indica vs Japonica closest to the root and 2) place the origin of the cBasmati lineage in close connection to both Japonica and cAus, either directly (Networks 1, 4 and 7) or through related immediate ancestry (Network 2).

The least likely studied networks (Networks 3 and 9) are at the same time those departing the most from scenarios proposed so far on the basis of similarity analyses. They display none of the features described above. Despite its intermediate ranking, Network 10 clearly has no biological pertinence: it displays Japonica as a late derivative of a hybridization between cAus and cBasmati, while it is now well-established that Japonica derives from the most ancient domestication in China. Networks 1, 7, 2 and 4 thus remain the scenarios of choice proposed by our method. Network 4 corresponds to the scenario that fits with most studies so far, because it conforms with the respective genetic distances between the three “pure” lineages, with Indica and cAus being closer to one another than with Japonica, and because cBasmati appears as a hybrid derivative of cAus x Japonica, which has been proposed by most recent studies. Yet this network is ranked at the forth position, just slightly better than Network 10.

Network 2, the third preferred scenario, proposes an ancient branch from a major Japonica-like foundation which combined with Indica to derive cAus and with Japonica to derive cBasmati. This is also compatible with the major hypotheses proposed so far by rice specialists, including the most recent one specifically focused on Basmati rices [80]. Network 7, which comes as the second best ranked scenario, keeps the hybrid origin of cBasmati as in Network 4, but indicates a recent divergence between Japonica and cAus. This would be surprising given the current and past geographical distributions of the three groups Japonica, Indica and cAus, which exhibits cAus in full vicinity with Indica.

Network 1 ranks far above the others. It is more complex, with two steps of hybridization. It features cBasmati as a hybrid derivative of the Japonica and cAus lineages, which conforms with current prominent hypotheses, but it also indicates a hybrid origin of the cAus lineage from early hybridization between Japonica and Indica lineages. This is not commonly mentioned as a possibility in the literature. According to our study, the respective contributions to cAus would be approximately 77:23 from Indica and Japonica (see Figure 6 in Supplementary Material), which concurs with the closer molecular proximity between Indica and cAus. Geographically, a cAus derivation from Indica x Japonica appears compatible with the current distribution of the three groups, with cAus concentrated in the Northeastern part of the Indian subcontinent where the southern silk road has established contacts between South Asia and East Asia since several centuries B.C. [81]. It would also explain a biological feature left unaccounted for so far: the wide cross compatibility of the cAus varieties which tends to make them fertile in crosses with both Indica and Japonica [82, 83].

We observe an agreement across analyses on networks 1, 7, 2 and 4: the proportion of Japonica present in cBasmati genome was estimated at approximately 73% (see Figure 6 in Supplementary Material). Although networks 7 and 4 are both *displayed* by networks 1 and 2, results on inheritance probabilities tend to link Network 1 to Network 4, and Network 2 to Network 7 (see Figure 6 in Supplementary Material).

Overall, Network 1 appears at the top of the ranking and thus can be taken as the first topology to be tested with more materials, including representatives of wild rice populations, and more finely focused analyses where ancestral recombination graphs can be tentatively sketched and assessed. For such analyses, SnappNet should benefit in the future from the new beast 2 package coupled MCMC [84], that tackles local optima issues thanks to heated chains.

## Discussion

In this paper, we introduced a new Bayesian method, SnappNet, dedicated to phylogenetic network inference. SnappNet has similar goals as `MCMCBiMarkers`, a method recently proposed by Zhu et al, but differs from this method in two main aspects. The first difference is due to the way the two methods handle the complexity of the sampled networks. Unlike binary trees that have a fixed number of branches given the number of considered species, network topologies can be of arbitrary complexity. Their complexity directly depends on the number of reticulations they contain. In MCMC processes, the complexity of sampled networks is regulated by the prior. `MCMCBiMarkers` uses descriptive priors: more precisely, it assumes a Poisson distribution for the number of reticulation nodes and an exponential distribution for the *diameter* of reticulation nodes. In contrast, SnappNet’s prior is based on that of Zhang et al., which explicitly relies on speciation and hybridization rates and is extendable to account for extinction and incomplete sampling [56].

In our simulation study, we investigated in detail the influence of these different priors. On two networks of moderate complexity (networks A and B), SnappNet and `MCMCBiMarkers` presented globally similar performances. Indeed, when we considered numbers of sites that are largely achieved in current phylogenomic studies (i.e. 10,000 or 100,000 sites), both methods were able to recover the true networks under this realistic framework. However, in presence of only a few sites (1,000 sites) which is unusual nowadays but still can be the case for poorly sequenced organisms, `MCMCBiMarkers` was found superior to SnappNet. On the other hand, when focusing on a more complex network (network C) containing reticulation nodes on top of one another, SnappNet clearly outperformed `MCMCBiMarkers` as soon as at least 10,000 sites were considered. SnappNet recovered the correct scenario in approximately 50% of cases whereas `MCMCBiMarkers` inferred this history in less than 5% of cases. In view of our simulation study, the birth hybridization process of [56] seems to be the most appropriate network prior for inferring complex evolution scenarios. This was expected since `MCMCBiMarkers`’s network prior, inspired by ancestral recombination graphs [85], penalize for networks with many reticulations (cf. [54]). However, complex scenarios are more and more frequent in the literature. A striking example is given by [86], who recently suggested a new thesis regarding the origin and evolution of *Citrus*. Cultivated varieties would result from hybridizations involving four ancestral taxons (*C. reticulata, C. medica, C. maxima, C. micrantha*). Some groups (e.g. the rough lemon) are the result of a direct hybridization between those ancestors whereas other groups are the product of a more complex history (e.g. the sweet orange). In this context, SnappNet appears to be a useful tool for researchers interested in evolutionary biology. To conclude the discussion on priors, we can also mention that, on simulated data, SnappNet’s accuracy did not really deteriorate with incorrect priors. The same behavior was observed by [2] for `MCMCBiMarkers`.

The second major difference between `MCMCBiMarkers` and SnappNet lies in the way they compute the likelihood of a network. This step is at the core of the bayesian analysis.

According to the authors of `MCMCBiMarkers`, this remains a major computational bottleneck and limits the applicability of their methods [61]. To understand the origin of this bottleneck, recall that the MCMC process of a Bayesian sampling explores a huge network space and that, at each exploration step, computing the likelihood is by far the most time consuming operation. Moreover, we need sometimes millions of runs before the chain converges. Thus, likelihood computation is a key factor on which to operate to be able to process large data sets.

The likelihood computation of `MCMCBiMarkers` consists in a bottom-up traversal, from the leaves to the root. Each time a reticulation node *r* is visited, each of the *n* lineages down this node have to be assigned to one of the two ancestors of *r*. The computational burden results from the fact that each of the 2^*n*^ possible partitions of the *n* lineages needs to be examined by `MCMCBiMarkers`. The split set of lineages will be merged only when reaching *lsa*(*r*), the *least stable ancestor* of, *r* that is the most recent node contained on all paths from the root to *r*. At this point, the “partial likelihoods” of all possible splits that were generated at *r* are combined to compute the partial likelihood at *lsa*(*r*). At each other reticulation *r*^*/*^ reached before *lsa*(*r*), the lineages again have to be split in two. As a result, the time required to compute the likelihood of a *blob* (i.e., a maximal biconnected subgraph [66]) grows exponentially with the number of reticulations it contains. More precisely, the time complexity of the likelihood computation in `MCMCBiMarkers` is in *O*(*sn*^4*k*+4^), where *k* is the *level* of the network and *s* is the size of the species network (i.e. its number of branches or its number of nodes) [2].

Similarly to `MCMCBiMarkers`, we compute the likelihood in a bottom-up process, and when reaching a reticulation node *r*, we also take into account the various ways lineages could have split. But the originality of SnappNet is to compute *joint conditional probabilities* for branches above a same reticulation node *r* (see Materials and methods section). The set of branches jointly considered increases when crossing other reticulation nodes in a same blob, but it can also decrease when crossing tree-nodes in the blob (i.e. nodes having one ancestor and several children). Of course, the time to compute each partial likelihood increases in proportion with the number of branches considered together. More precisely, SnappNet runs in *O*(*sn*^*2k′+2)*^), where *k*^*′*^ is the maximum number of branches simultaneously considered in a partial likelihood. The interest in depending on *k*^*′*^ instead of *k* (the number of reticulations in a blob), is that for some blobs, we can resort to a bottom-up traversal of the blob that limits *k*^*/*^ to a small constant and process the blob in polynomial time in *n*, while `MCMCBiMarkers` will still requires an exponential time in *k*.

Our results from simulated data conforms to the above theoretical discussion. For a single likelihood evaluation, SnappNet was found to be largely faster than `MCMCBiMarkers` on networks containing reticulation nodes on top of one another. Besides, SnappNet required substantially less memory than `MCMCBiMarkers`. This speed gain enables us to consider complex evolution scenarios in our Bayesian analyses. Unfortunately, on rice real data, although many complex networks were evaluated, the convergence of the Markov Chain was not achieved in a reasonable time. We could have reduced the number of sampled lineages for proper mixing as in [56], but we finally adopted a penalized likelihood method, keeping the same original data sets. Recall that in the penalized likelihood framework, the imposed penalty can be viewed as the prior in a Bayesian study. Network 1, containing reticulation nodes on top of one another, was found as the best explanation of the rice evolution.

In future research, this scenario will be studied more thoroughly, by considering more rice varieties and by adding also wild rice populations, which will allow a more detailed comparison with the existing histories present in the literature (e.g. [14, 75, 80, 87]). There is today a large debate on whether a unique domestication [75, 80] or 3 separate domestications [14] took place. By a deep understanding of the domestication process, it becomes possible for geneticists to reintroduce some wild diversity into cultivars.

As highlighted in [2], polymorphic sites are helpful for recovering networks. However, when only these sites are taken into account, it is essential to condition for the fact that invariant sites have been discarded. Indeed, on simulated data, we observed that without the correction factor, the estimated network height and the estimated network length were largely biased. This behavior was already observed in Snapp by [1], in a species tree context. Rice real data have been analyzed accordingly. In the future, in order to handle more sites in practice, we should extend the MSNC model to allow recombination events between loci. Recall that we have limited our rice study to 12,000 markers sampled along the genome because the model assumes independence between sampled sites, as does also Snapp’s model, from which we inherit. As mentioned in the review of [41], in order to model recombination properly, the study of networks within networks is an area for future research. A possibility would be to exploit existing results on Ancestral Recombination Graphs (see for instance [88]).

Last, it would be of high interest to study the influence of ILS on SnappNet’s ability to recover networks. As in [89], we noticed on simulated data that ILS helps to recover networks. For instance, on the complex network C (Fig. 6), the branch located below the hybridization node *H*_2_, was calibrated in order to observe some ILS on that branch. Indeed, in a preliminary analysis with a longer branch, SnappNet was not able to recover the true network. In other words, the statistical learning is facilitated by several lineages entering the reticulation node and choosing different paths, as theorized by [44]. This aspect should be investigated in more details in the future.

To conclude, a large amount of methodological questions on phylogenetic networks remain open. At the end of their paper, the authors of `MCMCBiMarkers` [2] concluded by mentioning that “An important direction for future research is improving the computational requirements of the method to scale up to data sets with many taxa”. Our present work is a first answer to this demand.

## Acknowledgments

We thank Wensheng Wang for kindly helping us choosing rice accessions for future rice comparative analyses and João D. Santos for providing a script of SNP extraction. This work was supported by the French Agence Nationale de la Recherche, “Investissements d’Avenir” program, through the 3 following projects: the project Genome Harvest (reference ID 1504-006, Labex Agro: ANR-10-LABX-0001-01), the project KIM Data & Life Sciences (I-SITE MUSE: ANR-16-IDEX-0006) and the project AdaptGrass (reference ID170544IA, I-SITE MUSE: ANR-16-IDEX-0006). It was also funded by the CGIAR Research Program on Rice Agrifood Systems (RICE). This study has been realized with the support of the ATGC platform of the French National Institute of Bioinformatics, with the support of the High Performance Computing Platform MESO@LR, financed by the Occitanie / Pyrénées-Méditerranée Region, Montpellier Mediterranean Metropole and the University of Montpellier. This work also benefited from the Montpellier Bioinformatics Biodiversity platform supported by the LabEx CeMEB, an ANR “Investissements d’avenir” program (ANR-10-LABX-04-01). Last, it was supported by the CIRAD - UMR AGAP HPC Data Center of the South Green Bioinformatics platform (http://www.south-green.fr/).

## Footnotes

1 the obtained tree, ((((Q,A),L),R),C), is just Network A where the branch having the smallest inheritance probability on top of a hybridization node is removed.

## References

1. Bryant D, Bouckaert R, Felsenstein J, Rosenberg NA, RoyChoudhury A. Inferring species trees directly from biallelic genetic markers: bypassing gene trees in a full coalescent analysis. Molecular biology and evolution. 2012;29(8):1917–1932.

2. Zhu J, Wen D, Yu Y, Meudt HM, Nakhleh L. Bayesian inference of phylogenetic networks from bi-allelic genetic markers. PLoS computational biology. 2018;14(1):e1005932.

3. Denoeud F, Carretero-Paulet L, Dereeper A, Droc G, Guyot R, Pietrella M, et al. The coffee genome provides insight into the convergent evolution of caffeine biosynthesis. science. 2014;345(6201):1181–1184.

4. Badouin H, Gouzy J, Grassa CJ, Murat F, Staton SE, Cottret L, et al. The sunflower genome provides insights into oil metabolism, flowering and Asterid evolution. Nature. 2017;546(7656):148.

5. Garsmeur O, Droc G, Antonise R, Grimwood J, Potier B, Aitken K, et al. A mosaic monoploid reference sequence for the highly complex genome of sugarcane. Nature communications. 2018;9(1):2638.

6. Cornillot E, Hadj-Kaddour K, Dassouli A, Noel B, Ranwez V, Vacherie B, et al. Sequencing of the smallest Apicomplexan genome from the human pathogen Babesia microti. Nucleic acids research. 2012;40(18):9102–9114.

7. Marra NJ, Stanhope MJ, Jue NK, Wang M, Sun Q, Bitar PP, et al. White shark genome reveals ancient elasmobranch adaptations associated with wound healing and the maintenance of genome stability. Proceedings of the National Academy of Sciences. 2019;116(10):4446–4455.

8. Consortium IH, et al. The international HapMap project. Nature. 2003;426(6968):789.

9. RGP. The 3,000 rice genomes project. GigaScience. 2014;3(1):2047–217X.

10. Bamshad MJ, Ng SB, Bigham AW, Tabor HK, Emond MJ, Nickerson DA, et al. Exome sequencing as a tool for Mendelian disease gene discovery. Nature Reviews Genetics. 2011;12(11):745.

11. Mansueto L, Fuentes RR, Chebotarov D, Borja FN, Detras J, Abriol-Santos JM, et al. SNP-Seek II: A resource for allele mining and analysis of big genomic data in Oryza sativa. Current Plant Biology. 2016;7:16–25.

12. Hernandez RD, Kelley JL, Elyashiv E, Melton SC, Auton A, McVean G, et al. Classic selective sweeps were rare in recent human evolution. science. 2011;331(6019):920–924.

13. Gravel S, Henn BM, Gutenkunst RN, Indap AR, Marth GT, Clark AG, et al. Demographic history and rare allele sharing among human populations. Proceedings of the National Academy of Sciences. 2011;108(29):11983–11988.

14. Civáň P, Craig H, Cox CJ, Brown TA. Three geographically separate domestications of Asian rice. Nature plants. 2015;1(11):15164.

15. Rouard M, Droc G, Martin G, Sardos J, Hueber Y, Guignon V, et al. Three new genome assemblies support a rapid radiation in Musa acuminata (wild banana). Genome biology and evolution. 2018;10(12):3129–3140.

16. Felsenstein J, Felenstein J. Inferring phylogenies. vol. 2. Sinauer associates Sunderland, MA; 2004.

17. Kingman JF. On the genealogy of large populations. Journal of applied probability. 1982;19(A):27–43.

18. Rannala B, Yang Z. Bayes estimation of species divergence times and ancestral population sizes using DNA sequences from multiple loci. Genetics. 2003;164(4):1645–1656.

19. Knowles LL, Kubatko LS. Estimating species trees: practical and theoretical aspects. John Wiley and Sons; 2011.

20. RoyChoudhury A, Felsenstein J, Thompson EA. A two-stage pruning algorithm for likelihood computation for a population tree. Genetics. 2008;.

21. Ebersberger I, Galgoczy P, Taudien S, Taenzer S, Platzer M, Von Haeseler A. Mapping human genetic ancestry. Molecular Biology and Evolution. 2007;24(10):2266–2276.

22. Degnan JH, Rosenberg NA. Gene tree discordance, phylogenetic inference and the multispecies coalescent. Trends in ecology & evolution. 2009;24(6):332–340.

23. Maddison WP. Gene Trees in Species Trees. Systematic Biology. 1997 09;46(3):523–536. Available from: https://doi.org/10.1093/sysbio/46.3.523.

24. Koonin EV, Makarova KS, Aravind L. Horizontal gene transfer in prokaryotes: quantification and classification. Annual Reviews in Microbiology. 2001;55(1):709–742.

25. Szöllósi GJ, Davín AA, Tannier E, Daubin V, Boussau B. Genome-scale phylogenetic analysis finds extensive gene transfer among fungi. Phil Trans R Soc B. 2015;370(1678):20140335.

26. Mallet J. Hybrid speciation. Nature. 2007;446(7133):279.

27. Morales L, Dujon B. Evolutionary role of interspecies hybridization and genetic exchanges in yeasts. Microbiology and Molecular Biology Reviews. 2012;76(4):721–739.

28. Glemin S, Scornavacca C, Dainat J, Burgarella C, Viader V, Ardisson M, et al. Pervasive hybridizations in the history of wheat relatives. bioRxiv. 2018;p. 300848.

29. Cui R, Schumer M, Kruesi K, Walter R, Andolfatto P, Rosenthal GG. Phylogenomics reveals extensive reticulate evolution in Xiphophorus fishes. Evolution. 2013;67(8):2166–2179.

30. Civáň P, Brown TA. Role of genetic introgression during the evolution of cultivated rice (Oryza sativa L.). BMC evolutionary biology. 2018;18(1):57.

31. Minamikawa MF, Nonaka K, Kaminuma E, Kajiya-Kanegae H, Onogi A, Goto S, et al. Genome-wide association study and genomic prediction in citrus: potential of genomics-assisted breeding for fruit quality traits. Scientific reports. 2017;7(1):4721.

32. Duranton M, Allal F, Fraïsse C, Bierne N, Bonhomme F, Gagnaire PA. The origin and remolding of genomic islands of differentiation in the European sea bass. Nature communications. 2018;9(1):2518.

33. Posada D, Crandall KA, Holmes EC. Recombination in evolutionary genomics. Annual Review of Genetics. 2002;36(1):75–97.

34. Bailes E, Gao F, Bibollet-Ruche F, Courgnaud V, Peeters M, Marx PA, et al. Hybrid origin of SIV in chimpanzees. Science. 2003;300(5626):1713–1713.

35. Huson DH, Rupp R, Scornavacca C. Phylogenetic networks: concepts, algorithms and applications. Cambridge University Press; 2010.

36. Nakhleh L. Evolutionary phylogenetic networks: models and issues. In: Problem solving handbook in computational biology and bioinformatics. Springer; 2010. p. 125–158.

37. Morrison DA. Introduction to Phylogenetic Networks. RJR Productions; 2011.

38. Baroni M, Semple C, Steel M. A framework for representing reticulate evolution. Annals of Combinatorics. 2005;8(4):391–408.

39. Hudson RR. Properties of a neutral allele model with intragenic recombination. Theoretical population biology. 1983;23(2):183–201.

40. Huson DH, Scornavacca C. A survey of combinatorial methods for phylogenetic networks. Genome biology and evolution. 2011;3:23–35.

41. Degnan JH. Modeling hybridization under the network multispecies coalescent. Systematic biology. 2018;67(5):786–799.

42. Fontaine MC, Pease JB, Steele A, Waterhouse RM, Neafsey DE, Sharakhov IV, et al. Extensive introgression in a malaria vector species complex revealed by phylogenomics. Science. 2015;347(6217):1258524.

43. Marcussen T, Sandve SR, Heier L, Spannagl M, Pfeifer M, Jakobsen KS, et al. Ancient hybridizations among the ancestral genomes of bread wheat. science. 2014;345(6194):1250092.

44. Zhu S, Degnan JH. Displayed trees do not determine distinguishability under the network multispecies coalescent. Systematic biology. 2016;66(2):283–298.

45. Huson DH, Scornavacca C. A Survey of Combinatorial Methods for Phylogenetic Networks. Genome Biology and Evolution. 2010 11;3:23–35. Available from: https://doi.org/10.1093/gbe/evq077.

46. Kubatko LS. Identifying hybridization events in the presence of coalescence via model selection. Systematic Biology. 2009;58(5):478–488.

47. Meng C, Kubatko LS. Detecting hybrid speciation in the presence of incomplete lineage sorting using gene tree incongruence: a model. Theoretical population biology. 2009;75(1):35–45.

48. Yu Y, Than C, Degnan JH, Nakhleh L. Coalescent histories on phylogenetic networks and detection of hybridization despite incomplete lineage sorting. Systematic Biology. 2011;60(2):138–149.

49. Yu Y, Degnan JH, Nakhleh L. The probability of a gene tree topology within a phylogenetic network with applications to hybridization detection. PLoS genetics. 2012;8(4):e1002660.

50. Yu Y, Ristic N, Nakhleh L; BioMed Central. Fast algorithms and heuristics for phylogenomics under ILS and hybridization. BMC bioinformatics. 2013;14(15):S6.

51. Yu Y, Dong J, Liu KJ, Nakhleh L. Maximum likelihood inference of reticulate evolutionary histories. Proceedings of the National Academy of Sciences. 2014;111(46):16448–16453.

52. Yu Y, Nakhleh L. A maximum pseudo-likelihood approach for phylogenetic networks. BMC genomics. 2015;16(10):S10.

53. Solís-Lemus C, Ané C. Inferring phylogenetic networks with maximum pseudolikelihood under incomplete lineage sorting. PLoS Genetics. 2016;12(3):e1005896.

54. Wen D, Yu Y, Nakhleh L. Bayesian inference of reticulate phylogenies under the multispecies network coalescent. PLoS genetics. 2016;12(5):e1006006.

55. Wen D, Nakhleh L. Coestimating reticulate phylogenies and gene trees from multilocus sequence data. Systematic biology. 2018;67(3):439–457.

56. Zhang C, Ogilvie HA, Drummond AJ, Stadler T. Bayesian inference of species networks from multilocus sequence data. Molecular biology and evolution. 2017;35(2):504–517.

57. Elworth RL, Ogilvie HA, Zhu J, Nakhleh L. Advances in computational methods for phylogenetic networks in the presence of hybridization. In: Bioinformatics and Phylogenetics. Springer; 2019. p. 317–360.

58. Bayzid MS, Warnow T. Naive binning improves phylogenomic analyses. Bioinformatics. 2013;29(18):2277–2284.

59. Bouckaert R, Heled J, Kühnert D, Vaughan T, Wu C, Xie D, et al. BEAST 2: a software platform for Bayesian evolutionary analysis. PLoS computational biology. 2014;10(4):e1003537.

60. Bouckaert R, Vaughan TG, Barido-Sottani J, Duchêne S, Fourment M, Gavryushkina A, et al. BEAST 2.5: An advanced software platform for Bayesian evolutionary analysis. PLoS computational biology. 2019;15(4):e1006650.

61. Zhu J, Nakhleh L. Inference of species phylogenies from bi-allelic markers using pseudo-likelihood. Bioinformatics. 2018;34(13):i376–i385.

62. Pardi F, Scornavacca C. Reconstructible phylogenetic networks: do not distinguish the indistinguishable. PLoS computational biology. 2015;11(4):e1004135.

63. Haldane J. The combination of linkage values and the calculation of distances between the loci of linked factors. J Genet. 1919;8(29):299–309.

64. Cavender JA. Taxonomy with confidence. Mathematical biosciences. 1978;40(3-4):271–280.

65. Cormen TH, Leiserson CE, Rivest RL, Stein C. Introduction to Algorithms, Third Edition. 3rd ed. The MIT Press; 2009.

66. Gambette P, Berry V, Paul C. The structure of level-k phylogenetic networks. In: Annual Symposium on Combinatorial Pattern Matching. Springer; 2009. p. 289–300.

67. Berry V, Scornavacca C, Weller M. Scanning Phylogenetic Networks is NP-hard. In: SOFSEM; 2020. In press.

68. Cardona G, Rosselló F, Valiente G. Extended Newick: it is time for a standard representation of phylogenetic networks. BMC bioinformatics. 2008;9(1):532.

69. Liu JS. Monte Carlo strategies in scientific computing. Springer Science & Business Media; 2008.

70. Rambaut A, Drummond AJ, Xie D, Baele G, Suchard MA. Posterior summarization in Bayesian phylogenetics using Tracer 1.7. Systematic biology. 2018;67(5):901–904.

71. Wang W, Mauleon R, Hu Z, Chebotarov D, Tai S, Wu Z, et al. Genomic variation in 3,010 diverse accessions of Asian cultivated rice. Nature. 2018;557(7703):43–49.

72. Akaike H. Information theory and an extension of the maximum likelihood principle. In: Selected papers of hirotugu akaike. Springer; 1998. p. 199–213.

73. Schwarz G, et al. Estimating the dimension of a model. The annals of statistics. 1978;6(2):461–464.

74. Glaszmann JC. Isozymes and classification of Asian rice varieties. Theoretical and Applied genetics. 1987;74(1):21–30.

75. Huang X, Kurata N, Wang ZX, Wang A, Zhao Q, Zhao Y, et al. A map of rice genome variation reveals the origin of cultivated rice. Nature. 2012;490(7421):497.

76. Civáň P, Ali S, Batista-Navarro R, Drosou K, Ihejieto C, Chakraborty D, et al. Origin of the aromatic group of cultivated rice (Oryza sativa L.) traced to the Indian subcontinent. Genome biology and evolution. 2019;11(3):832–843.

77. Santos JD, Chebotarov D, McNally KL, Bartholomé J, Droc G, Billot C, et al. Fine scale genomic signals of admixture and alien introgression among Asian rice landraces. Genome biology and evolution. 2019;11(5):1358–1373.

78. Drummond AJ, Bouckaert RR. Bayesian evolutionary analysis with BEAST. Cambridge University Press; 2015.

79. Larget B. The estimation of tree posterior probabilities using conditional clade probability distributions. Systematic biology. 2013;62(4):501–511.

80. Choi JY, Platts AE, Fuller DQ, Wing RA, Purugganan MD, et al. The rice paradox: multiple origins but single domestication in Asian rice. Molecular biology and evolution. 2017;34(4):969–979.

81. Chakraborty D, Ray A. Population genetics analyses of North-East Indian indigenous rice landraces revealed divergent history and alternate origin of aroma in aus group. Plant Genetic Resources. 2019;17(5):437–447.

82. Morinaga T, Kuriyama H. Japonica type rice in the subcontinent of India and Java. Japanese Journal of Breeding. 1955;5(3):149–153.

83. Morinaga T, Kuriyama H. Intermediate type of rice in the subcontinent of India and Java. Japanese Journal of Breeding. 1958;7(4):253–259.

84. Mueller NF, Bouckaert R. Adaptive parallel tempering for BEAST 2. bioRxiv. 2020;p. 603514.

85. Bloomquist EW, Suchard MA. Unifying vertical and nonvertical evolution: a stochastic ARG-based framework. Systematic biology. 2010;59(1):27–41.

86. Wu GA, Terol J, Ibanez V, López-García A, Pérez-Román E, Borredá C, et al. Genomics of the origin and evolution of Citrus. Nature. 2018;554(7692):311.

87. Zhao Q, Feng Q, Lu H, Li Y, Wang A, Tian Q, et al. Pan-genome analysis highlights the extent of genomic variation in cultivated and wild rice. Nature genetics. 2018;50(2):278.

88. Gusfield D. ReCombinatorics: the algorithmics of ancestral recombination graphs and explicit phylogenetic networks. MIT press; 2014.

89. Zhu S, Degnan JH. Displayed trees do not determine distinguishability under the network multispecies coalescent. Systematic biology. 2017;66(2):283–298.

